# Molecular mechanisms of E-Syt-mediated stress resistance

**DOI:** 10.64898/2026.06.23.733366

**Authors:** Francisco Benitez-Fuente, Javier Collado, Jorge Morello-López, Raquel Pagano-Marquez, Noemi Ruiz-Lopez, Jenny Keller, Miguel A. Botella, Rubén Fernández-Busnadiego

## Abstract

Membrane contact sites (MCS) between the endoplasmic reticulum (ER) and the plasma membrane (PM) enable direct intermembrane exchange of signals and metabolites. The Extended Synaptotagmins (E-Syts) are an evolutionary conserved family of ER-PM tethers essential to maintain PM integrity under stress conditions. To investigate the underlying molecular mechanisms, we employed cellular reconstitution experiments in yeast and plants. We show that E-Syt-mediated stress tolerance relies on ER-PM MCS targeting, which requires the E-Syt N-terminal membrane anchor, a minimal set of two C2 domains and an SMP domain. C2 domains are sufficiently conserved that interspecies domains can sustain both PM localization and stress response. The role of the SMP domain in ER-PM localization is also conserved, but SMP function in stress resistance is species-specific. Furthermore, cryo-electron tomography uncovers a scaffolding role for the SMP domain in maintaining ER-PM distance, and in the formation of ER membrane peaks with extreme curvature that appear necessary for stress tolerance. Collectively, our findings reveal the individual and synergistic roles of all E-Syt modules in maintaining cellular homeostasis under stress.

## Introduction

Membrane contact sites (MCS) between the endoplasmic reticulum (ER) and the plasma membrane (PM), where both membranes come within 30 nm of each other^1^, provide essential platforms for regulation of calcium and lipid homeostasis^2,3^, cell signaling pathways^4,5^ and organelle biogenesis^6^. These functions are mediated by proteins that often also act as tethers, structurally bridging the membranes^7^. Extended Synaptotagmins (E-Syts)^8^ are a conserved family of ER-PM tethers, with homologs from human (hE-Syts) to yeast (Tricalbins; Tcbs) and plants (Synaptotagmins; SYTs) (Figure 1A). E-Syts can connect the ER to the PM by simultaneously anchoring to ER tubules^9–12^ through their N-terminal domain, and docking to the PM via their C-terminal C2 domains^7,10,13–15^. Their central module is a synaptotagmin-like mitochondrial-lipid–binding protein (SMP) domain, which can be found in several MCS-resident proteins involved in lipid transport^16–19^. Previous findings show that E-Syt SMP domains can bind and transfer lipids^13,19–22^, and mediate E-Syts homo/heterodimerization^9,20,23–28^ (Figure S1E).

**Figure 1:**
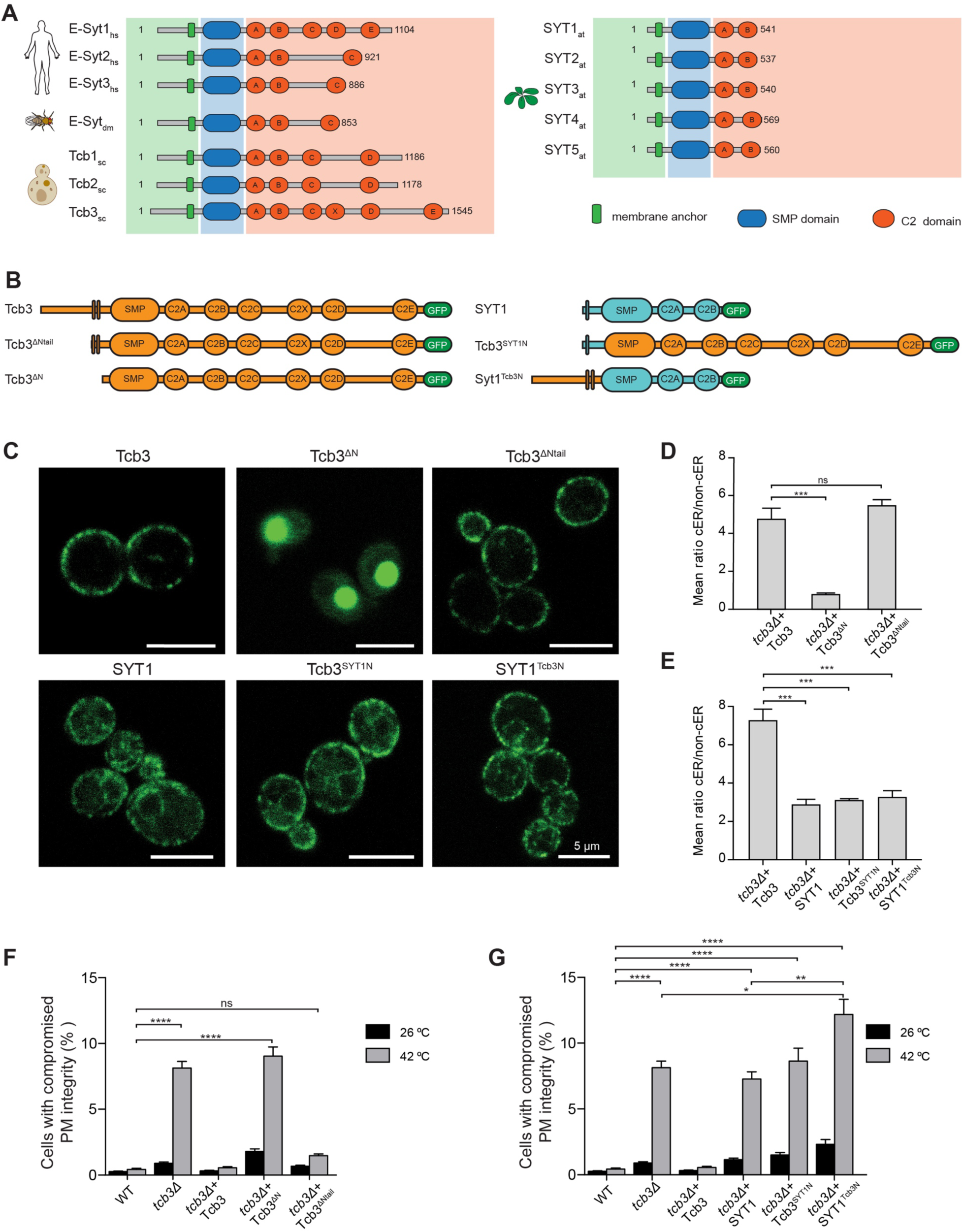
The E-Syt N-terminal module is essential for ER-PM MCS targeting in yeast. **A**: Schematic depiction of E-Syt modules across different species. hs: *Homo sapiens*, dm: *Drosophila melanogaster*, at: *Arabidopsis thaliana*, sc: *Saccharomyces cerevisiae*. Originally, only five C2 domains were annotated for Tcb3 but AlphaFold predicts an additional domain between C2C and C2D, which we refer to as C2X (See also Figure S1B). **B**: Schematic representation of constructs used in C-G. **C**: Subcellular localization of the constructs monitored by confocal imaging. **D**: Quantification of cER localization of Tcb3 truncation mutants. **E**: Quantification of cER localization of Tcb3-SYT1 chimeras. In D and E, the localization was asserted by the ratio between cortical (cER) and non-cortical (non-cER) fluorescence. See also Figure S2 and Table S2. **F**: PM integrity assay of *tcb3*Δ cells complemented with Tcb3 full length or Tcb3 truncation mutants upon 10 min incubation at 42 °C. **G**: PM integrity assay of *tcb3*Δ cells complemented with Tcb3 full length, SYT1 full length or Tcb3-SYT1 chimeras upon 10 min incubation at 42 °C. In F and G, the Y axis shows the % of cells with intracellular PI signal in flow cytometry experiments. The plots in D-G show average values for each condition + SEM (error bars). ns, *, **, *** and **** indicate p-value > 0.05, p-value < 0.05, p-value < 0.01, p-value < 0.001 and p-value < 0.0001, respectively, using one-way (D, E) and two-way (F, G) Anova followed by Tukey’s multiple comparison tests. In F and G, significance is only displayed for selected comparisons. P-values for all comparisons can be found in Table S1.

Despite their shared domain organization, E-Syt family members differ substantially. Experimental data^9^ and computational predictions suggest that in *Saccharomyces cerevisiae*, *Drosophila melanogaster* (hereafter, *Drosophila*) and humans, the N-terminal module most likely consists of a cytosolic unstructured tail followed by a hairpin structure, which inserts its central part in the ER membrane (Figure S1A). In contrast, in *Arabidopsis thaliana* (hereafter, *Arabidopsis*) the SYT1 N-terminus likely folds in a single transmembrane helix^29^ (Figure S1A), suggesting major differences in how the various E-Syts are targeted to ER-PM MCS. E-Syts also possess a variable number of C2 domains ranging from two in *Arabidopsis* SYT1, up to six C2 domains in *S. cerevisiae* Tcb3 (Figure1A, Figure S1B), which might reflect differences in establishment and/or regulation of ER-PM MCS. It has been proposed that some C2 domains bind the PM constitutively^7,10,13–15,30,31^, most likely via a conserved positively charged patch on their surface (Figure S1C, red circles), while others may bind the PM in Ca^2+^ -dependent manner. Indeed, SYT1, dmE-Syt^15^ and hE-Syts^13,32,33^ possess at least one Ca^2+^-binding C2 domain, which strengthens E-Syt ER-PM MCS localization in response to Ca^2+^ signals^15,30,32,33^.

E-Syts have emerged as important stress resistance factors. *Arabidopsis* SYT1 loss of function mutant has no detectable phenotype in normal conditions, but shows strong hypersensitivity to abiotic stresses, such as freezing, salt and mechanical stress^14,19,22,31,34^. Similarly, yeast cells depleted of Tcbs display no obvious basal phenotypes, but suffer increased PM damage upon heat stress^10^. *Drosophila* photoreceptors also appear normal in dmE-Syt knock-out under standard light conditions, but undergo degeneration under constant illumination^35^. E-Syt knock-out mice show no effects on animal development or viability^36–38^, but compromised embryonic fibroblast migration and survival was observed under stress^36^. Taken together, these data point towards an important role of the E-Syt family in stress response. However, the underlying molecular mechanisms are poorly understood, partly due to the complex E-Syt domain organization, as well as its evolutionary variability.

Here, we sought out to decipher the role of E-Syts domains in ER-PM MCS targeting and stress tolerance, combining functional studies, light microscopy and cryo-electron tomography (cryo-ET). To this end, we focused on yeast *S. cerevisiae* Tricalbin 3 (Tcb3) and *Arabidopsis* SYT1, as their respective mutants exhibit the most penetrant phenotypes in yeast and plants, respectively. We used truncation mutants and protein chimeras to explore the structure-function relationship of all E-Syt domains. Our results clarify the synergistic roles of the different E-Syt modules across evolution.

## Results

### The ER membrane anchor is necessary but not sufficient to target E-Syts to ER-PM MCS

First, we analyzed the molecular determinants of E-Syt ER-PM MCS localization and stress tolerance using yeast as model system. In contrast to plants, where only 5-10 % of the PM is involved in ER-PM MCS^36^, in *S. cerevisiae* more than a third of the PM surface is engaged in MCS with the ER^7,40,41^, defining the ER region participating in these contacts as “cortical ER” (cER). Thus, targeting of proteins to ER-PM MCS can be easily monitored by measuring their peripheral localization within the cER by fluorescence imaging^7,9,11^ (Figure 1C). To study stress tolerance, we incubated cells with membrane-impermeable propidium iodide (PI), whose intracellular accumulation reflects the loss of PM integrity^40,42^.

In yeast, Tcb3 localizes to the cER, and cells lacking Tcb3 (*tcb3Δ*) display PM integrity defects comparable to cells lacking all Tcbs^10,12,40^. To understand which regions of Tcb3 mediate these phenomena, we reintroduced various Tcb3-GFP constructs in *tcb3Δ* cells (Figure 1B). First, we confirmed that a full-length Tcb3 construct localized cortically (Figure 1C, D, Figure S2) and rescued the *tcb3Δ* PM integrity defect (Figure 1F). As expected, a Tcb3 construct lacking the complete N-terminus, including the ER membrane anchor (Tcb3^ΔN^, Figure 1B), resulted in a total loss of ER binding and seemed to accumulate in the vacuole (Figure 1C, D). Accordingly, this construct resulted in PM integrity defects comparable to *tcb3Δ* cells (Figure 1F). In contrast, a Tcb3 construct lacking only the unstructured tail N-terminal to the membrane anchor (Tcb3^ΔNtail^, Figure 1B, Figure S1A) localized to the cER similarly to the full-length protein (Figure 1C, D) and rescued the *tcb3*Δ phenotype to levels similar to wild type (WT) Tcb3 (Figure 1F). Thus, while ER membrane anchoring is fully required for Tcb3 localization and stress tolerance function, the long N-terminal tail does not seem to play a major role in these phenomena.

In contrast to the hairpin structure found in Tcb3 and metazoan E-Syts, *Arabidopsis* SYT1 may use a single transmembrane helix^29^ as ER anchor (Figure S1A). To test the role of the hairpin in cER targeting, we expressed full-length SYT1 in yeast (Figure 1B). Contrary to the exclusive ER-MCS localization of Tcb3 in yeast, and SYT1 in plants^31^, SYT1 was distributed throughout the entire ER network in yeast (Figure 1C, E). This suggested that the transmembrane domain of SYT1, while still being able to anchor the protein to the yeast ER, cannot localize SYT1 specifically to the cER. Accordingly, upon heat stress, *tcb3*Δ cells expressing full length SYT1 exhibited PM damage similar to *tcb3*Δ cells (Figure 1G).

As the SMP and C2 domains are also required to target Tcb3 to ER-PM MCS^7,9,10,16^, we asked whether substituting only the Tcb3 N-terminus by the transmembrane domain of SYT1 (Tcb3^SYT1N^, Figure 1B) would suffice to rescue the cER localization and the *tcb3*Δ phenotype. However, similarly to full length SYT1, Tcb3^SYT1N^ localized throughout the entire ER (Figure 1C, E) and did not rescue the *tcb3*Δ phenotype (Figure 1G). This implies that, to confer heat-stress tolerance, Tcb3 must exclusively localize to the cER, and this process absolutely requires the N-terminal hairpin anchor. However, the hairpin alone is not sufficient, as cells expressing a construct of SYT1 in which the transmembrane domain was replaced by that of Tcb3 (SYT1^Tcb3N^, Figure 1B) showed a general ER localization (Figure 1C, E) and compromised PM integrity (Figure 1G). In fact, the phenotype was even more severe than that of *tcb3*Δ, possibly hinting to a role of the membrane anchor in protein-protein interactions. Collectively, these results demonstrate that while the ER membrane anchor is essential for E-Syts targeting to ER-PM MCS, additional domains and/or protein interactions are needed for proper cER localization.

### ER-PM MCS targeting by the C-terminal C2 domains is a conserved feature of E-Syts

In agreement with the key roles of E-Syt C2 domains in PM binding, the deletion of all six C2 domains (Tcb3^ΔC2^) disrupts cER localization^7,10,16^, and, as expected^10^, resulted in PM damage upon heat shock (Figure 2E). To explore the function of the various E-Syt C2 domains in more detail, we tested combinations of Tcb3 C2 domain deletions (Figure 2A). We sequentially deleted the three N-terminal or the three C-terminal C2 domains of Tcb3 (Tcb3^ΔC2ABC^ and Tcb3^ΔC2XDE^, respectively; Figure 2A). Tcb3^ΔC2ABC^ localized to the cER, similarly to full length Tcb3 (Figure 2B, C), and showed a significant but mild increased hypersensitivity to heat-stress (Figure 2E). On the other hand, Tcb3^ΔC2XDE^ showed an impaired cER localization and a compromised PM integrity similar to *tcb3Δ* (Figure 2B, C and E). These results indicate that the three C-terminal C2 domains (C2XDE) are sufficient to direct Tcb3 to the cER and can largely sustain its heat-tolerance role, while the N-terminal C2 domains (C2ABC) are dispensable for cER localization and play a limited role in Tcb3 stress-resistance function.

**Figure 2:**
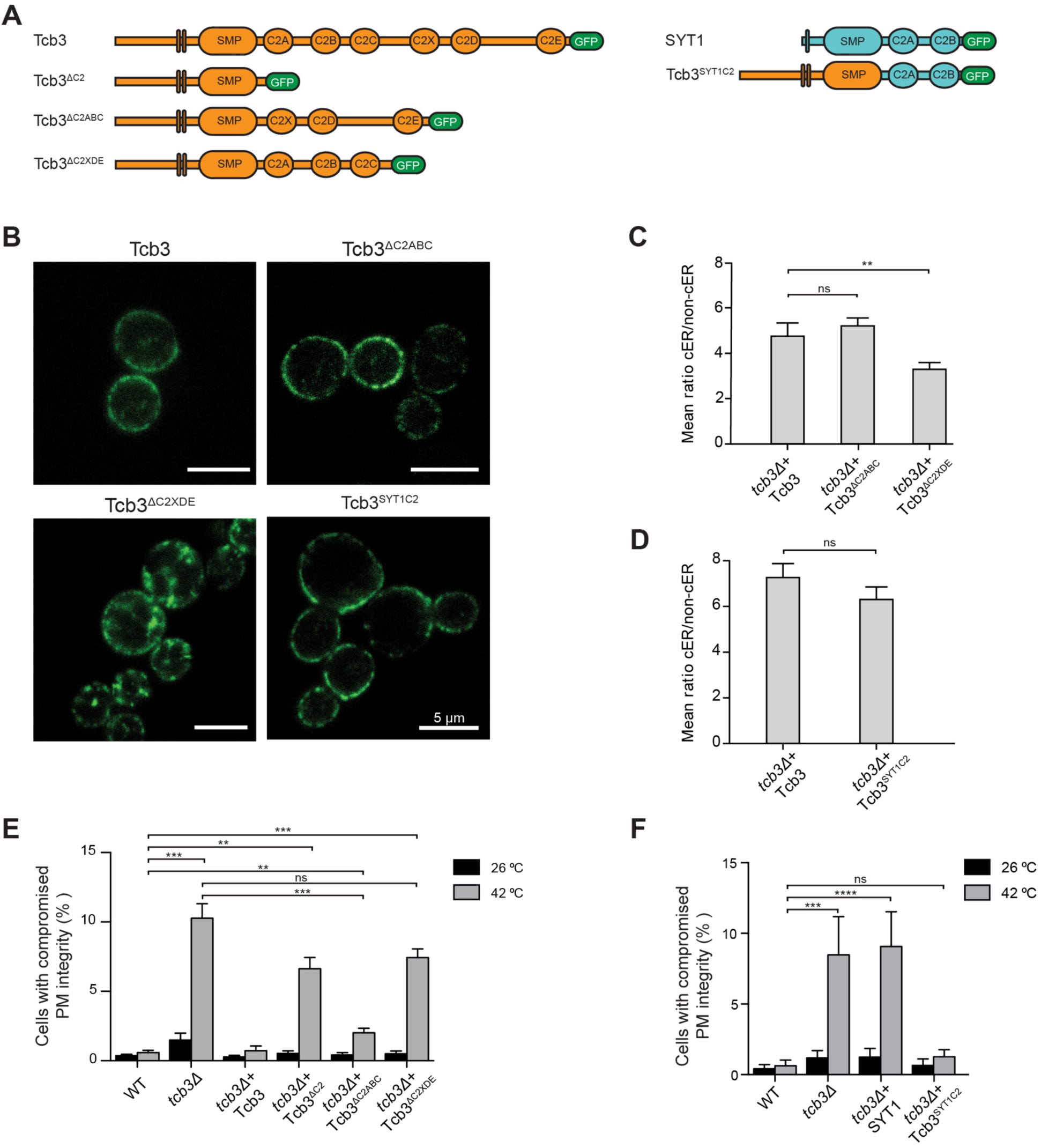
The C-terminal C2 domains are necessary to target E-Syts to yeast ER-PM MCS. **A**: Schematic representation of constructs used in B-F. **B**: Subcellular localization of the constructs monitored by confocal imaging. **C**: Quantification of cER localization of Tcb3 truncation mutants. **D**: Quantification of cER localization of Tcb3-SYT1 chimeras. In C and D, the localization was asserted by the ratio between cortical (cER) and non-cortical (non-cER) fluorescence. See also Figure S2 and Table S2. **E**: PM integrity assay of *tcb3*Δ cells complemented with Tcb3 full length or Tcb3 truncation mutants upon 10 min incubation at 42 °C. **F**: PM integrity assay of *tcb3*Δ cells complemented with Tcb3 full length, SYT1 full length or the Tcb3-SYT1 chimera upon 10 min incubation at 42 °C. In E and F, the Y axis shows the % of cells with intracellular PI signal in flow cytometry experiments. The plots C-F show average values for each condition + SEM (error bars). ns, **, *** and **** indicate p-value > 0.05, p-value < 0.01, p-value < 0.001 and p-value < 0.0001 respectively, using one-way (C, D) and two-way (E, F) Anova followed by Tukey’s multiple comparison tests. In E and F, significance is only displayed for selected comparisons. P-values for all comparisons can be found in Table S1.

This made us wonder to what extent interspecies C2 domains could functionally substitute those of Tcb3. To address this question, we expressed a construct where all six C2 domains of Tcb3 were replaced by the two C2 domains of SYT1 (Tcb3^SYT1C2^, Figure 2A). Interestingly, Tcb3^SYT1C2^ localization was indistinguishable from that of full length Tcb3 (Figure 2B, D), and the construct fully rescued the *tcb3*Δ PM integrity defect (Figure 2F). Thus, the two SYT1 C2 domains appear functionally equivalent to the whole set of six C2 domains of Tcb3 in these assays. Taken together, these results suggest that the mechanisms driving PM targeting and stress-sensing by C2 domains are evolutionary conserved.

### The SMP domain targets E-Syts to ER-PM MCS independently of its role in stress tolerance

So far, our results suggest that, for proper localization at ER-PM MCS in yeast, Tcb3 needs both its N-terminal hairpin anchor and a minimum set of two C2 domains, the latter being interchangeable among species. However, current evidence suggests that the SMP domain is also involved in targeting E-Syts to ER-PM MCS ^7,10,16^. Consistently, a Tcb3 construct lacking the SMP domain (Tcb3 ^ΔSMP^, Figure 3A) was not only unable to properly localize at the cER^10^ (Figure 3B, C) but also to maintain PM integrity in yeast^10^ (Figure 3D).

**Figure 3:**
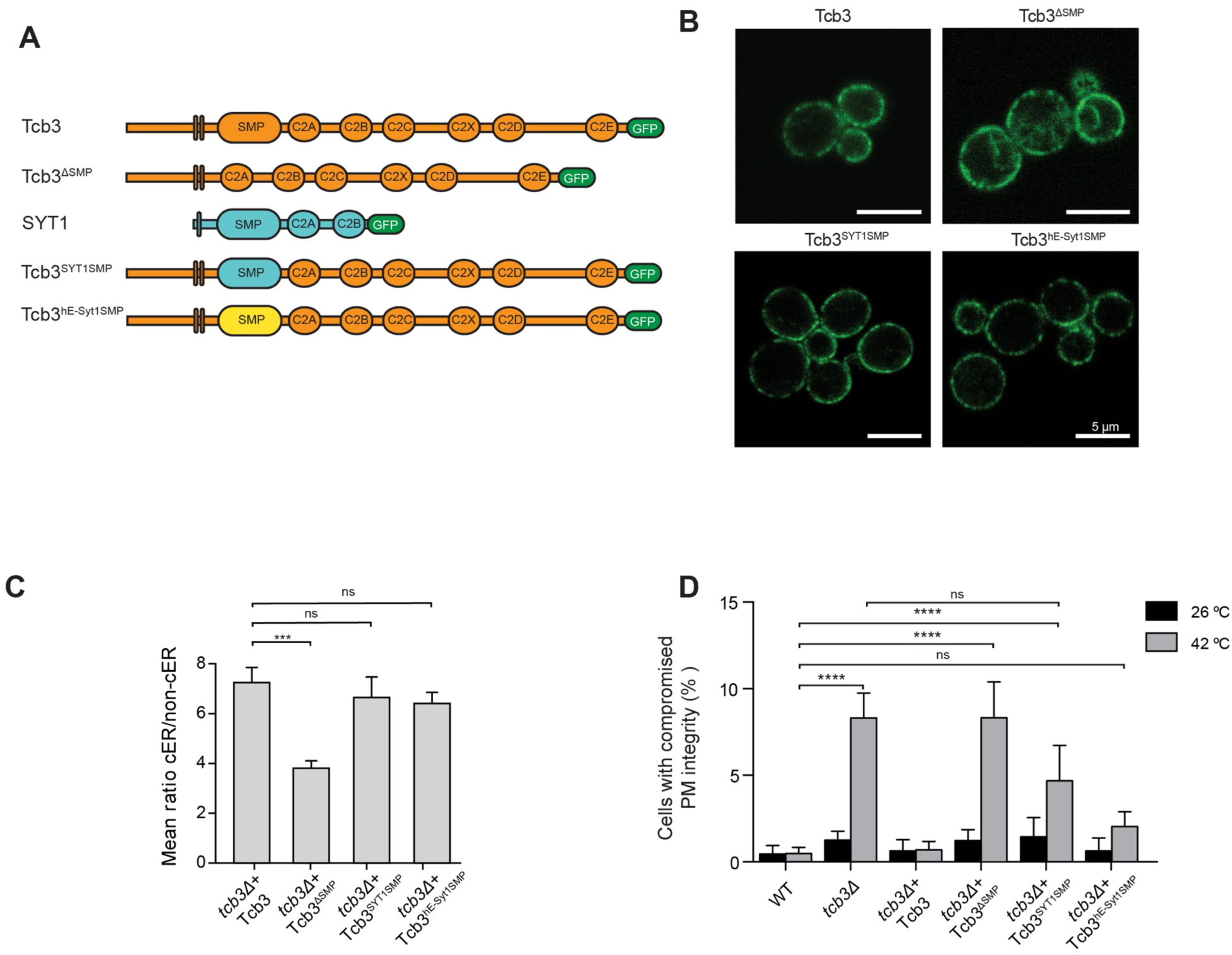
The SMP domain is required to target E-Syts to yeast ER-PM MCS. **A**: Schematic representation of constructs used in B-D. **B**: Subcellular localization of the constructs monitored by confocal imaging. **C**: Quantification of cER localization of the constructs. Localization was asserted by the ratio between cortical (cER) and non-cortical (non-cER) fluorescence. See also Figure S2 and Table S2. **D**: PM integrity assay of *tcb3*Δ cells complemented with Tcb3 full length, Tcb3 truncation mutant, SYT1 full length, Tcb3-SYT1 or Tcb3-hE-Syt1 chimeras upon 10 min incubation at 42 °C. The Y axis shows the % of cells with intracellular PI signal in flow cytometry experiments. The plots in C and D show average values for each condition + SEM (error bars). ns, *** and **** indicate p-value > 0.05, p-value < 0.001 and p-value < 0.0001, respectively, using one-way (C) and two-way (D) Anova followed by Tukey’s multiple comparison tests. In D, significance is only displayed for selected comparisons. P-values for all comparisons can be found in Table S1.

To gain further insight into the ER-PM MCS targeting role of the SMP domain, we generated Tcb3 variants in which the SMP domain was replaced by that of *Arabidopsis* SYT1 (Tcb3^SYT1SMP^, Figure 3A) or human hE-Syt1 (Tcb3^hE-Syt1SMP^, Figure 3A). Surprisingly, both Tcb3^SYT1SMP^ and Tcb3^hE-Syt1SMP^ exhibited a subcellular localization undistinguishable from that of Tcb3 (Figure 3B, C), suggesting that, similarly to C2 domains, cER targeting by SMP domains is also conserved across the E-Syt family. We next ascertained the ability of Tcb3^SYT1SMP^ and Tcb3^hE-Syt1SMP^ to sustain PM integrity upon heat shock. Intriguingly, while Tcb3^hE-Syt1SMP^ largely rescued the heat shock sensitivity of *tcb3*Δ, Tcb3^SYT1SMP^ resulted in significant PM integrity defects (Figure 3D). Collectively, these results indicate that the SMP domain plays a conserved structural role in targeting E-Syts at ER-PM MCS. The fact that Tcb3^SYT1SMP^ showed cER localization but no functional rescue indicates that the role of the E-Syt SMP domain in localization and stress-tolerance are differently regulated.

### Functional specificity of the SMP domain in *Arabidopsis*

To further characterize the roles of the SMP domain, we performed functional and localization experiments in plants. To visualize ER-PM MCS in plants, we focused on the cortical region of epidermal cells on the abaxial side of the leaf, where mouth-shaped stomata are also visible (Figure 4A). We used transient transformation of *Nicotiana benthamiana* (tobacco) expressing fluorescently-tagged SYT1 constructs (Figure 4B), as well as stable transgenic *Arabidopsis* plants expressing these constructs in a loss-of-function *syt1* background^14,31^. Using confocal microscopy, we determined the localization of these constructs and compared them to that of WT SYT1 and MAPPER, an artificial ER-PM tether that localizes at ER-PM MCS^19,31,43,44^ (Figure S3A, Figure 4C). We analyzed SYT1 constructs in which the SMP domain was swapped with that of yeast Tcb3 (SYT1^Tcb3SMP^, Figure 4B) or hE-Syt1 (SYT1^hE-Syt1SMP^, Figure 4B), both of which localized at ER-PM MCS similarly to SYT1 and MAPPER (Figure S3A, Figure 4C). These results reinforce the idea that the role of the SMP domain in ER-PM localization is conserved across species.

**Figure 4:**
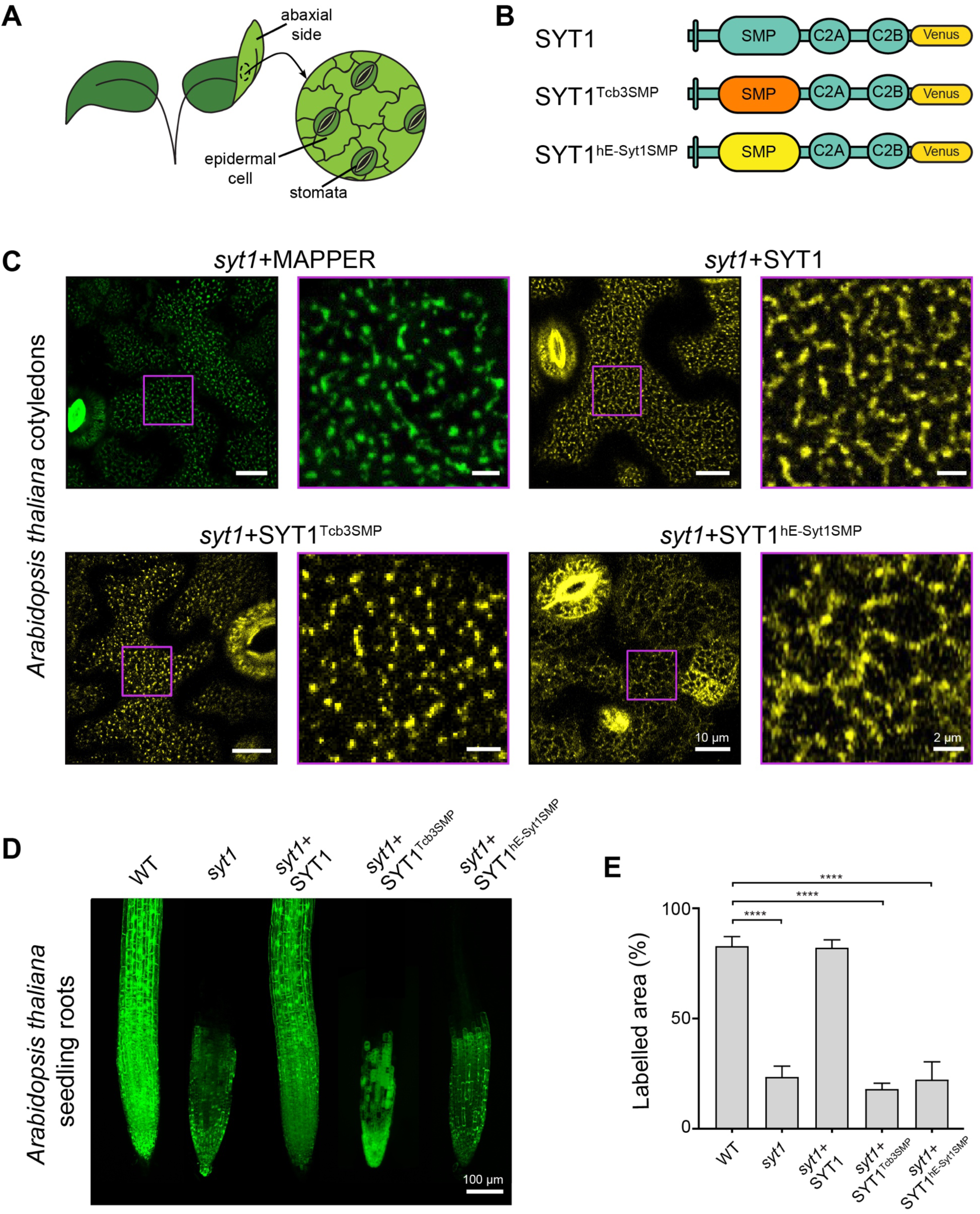
Localization and stress-resistance roles of the E-Syt SMP domain in Arabidopsis. **A:** Scheme depicting the region of the leaf used for localization experiments. **B:** Schematic representation of constructs used in C-E. **C:** Subcellular localization of MAPPER and SYT1 constructs depicted in B, at the cortical region of epidermal cells of stably transformed *Arabidopsis* cotyledons in *syt1* loss-of-function mutant background by confocal imaging. **D:** Confocal microscope images of cold-treated 6-days old *Arabidopsis* seedling roots stained with fluorescein diacetate. **E:** Quantification of D. The percentage of fluorescent area in different square ROIs along the root was measured. The bar plot displays means + SE (error bars). **** indicates p-value < 0.0001 using one-way Anova followed by Tukey’s multiple comparison test. Significance is asserted in comparison to WT and only displayed for selected comparisons. P-values for all comparisons can be found in Table S3.

We next evaluated the functionality of these constructs on stress-tolerance, exploiting the key role of SYT1 in resistance to cold stress in plants^14,19^. We stained seedling roots with fluorescein diacetate, which becomes fluorescent after hydrolysis by esterases only in living cells^14,19^. In agreement with previous work^19^, *syt1* mutant roots displayed a large darker region, indicating cell disruption, which was absent in WT (Figure 4D, E). As expected^19^, re-expression WT SYT1 fully rescued the tolerance to cold stress to WT levels (Figure 4D, E). In contrast, the number of fluorescent cells in the roots of SYT1^Tcb3SMP^ and SYT1^hE-Syt1SMP^ seedlings was similar to *syt1*, indicating that neither the hE-Syt1 nor Tcb3 SMP domain could fulfill the stress tolerance role of the SYT1 SMP domain in plants (Figure 4D, E). Taken together, these results are consistent with our findings in yeast, pointing to two different roles for the SMP domain of E-Syts: 1) a conserved role ensuring ER-PM MCS localization, and 2) a species-specific functional role required for PM integrity during stress episodes.

### Structural correlates of Tcb3 function in yeast ER-PM MCS

Cryo-ET enables high-resolution imaging of cellular landscapes in near-native conditions^45^, and is therefore well-suited to visualize the fine morphology of MCS^1,10,12,46–48^. Our previous cryo-ET study of ER-PM MCS in yeast showed that Tcb3 is involved in the formation of regions of high curvature at the cER (“cER peaks”), which may facilitate lipid transfer between the ER and the PM to maintain PM integrity^10^ under stress. Although these peaks were also observed under basal conditions, they were significantly upregulated upon heat shock^10^.

Therefore, here we analyzed ER-PM MCS of selected Tcb3 constructs in yeast upon heat shock using cryo-ET (Figure 5). In line with our previous results^10^, we observed peaks in WT cells (Figure 5A, blue arrowheads, Figure 5H), along with pleiomorphic tethering densities extending from the sides of the cER peaks to the PM (Figure 5A, black arrowheads). No peaks were detected in *tcb3*Δ cells (Figure 5B, H), but peak formation was recovered by re-expressing Tcb3 full length (Figure 5C, blue arrowheads, Figure 5H), confirming the direct involvement of Tcb3 in cER peak establishment. In contrast, no cER peaks were observed in *tcb3*Δ cells expressing Tcb3^ΔSMP^ (Figure 5D, H), in line with the defective localization and no functional rescue observed for this construct^10^ (Figure 3B-D). Next, we examined the capacity of Tcb3^SYT1SMP^ and Tcb3^hE-Syt1SMP^ to induce cER peaks, as both constructs correctly localized to cER (Figure 3B, C) but only Tcb3^hE-Syt1SMP^ provided significant functional rescue (Figure 3D). Interestingly, whereas cER peaks were observed in both conditions (Figure 5E, F blue arrowheads, Figure 5H), Tcb3^hE-Syt1SMP^ led to WT-like levels while a very strong reduction was observed for Tcb3^SYT1SMP^ (Figure 5H), in line with their difference in functional rescue. Lastly, Tcb3^SYT1C2^, which displayed proper cER localization and functional rescue (Figure 2B, D and F), promoted cER peak formation at levels similar to WT (Figure 5G blue arrowheads, Figure 5H). Collectively, these results establish a direct correlation between the SMP function in maintenance of PM integrity and its role in the formation of cER peaks.

**Figure 5:**
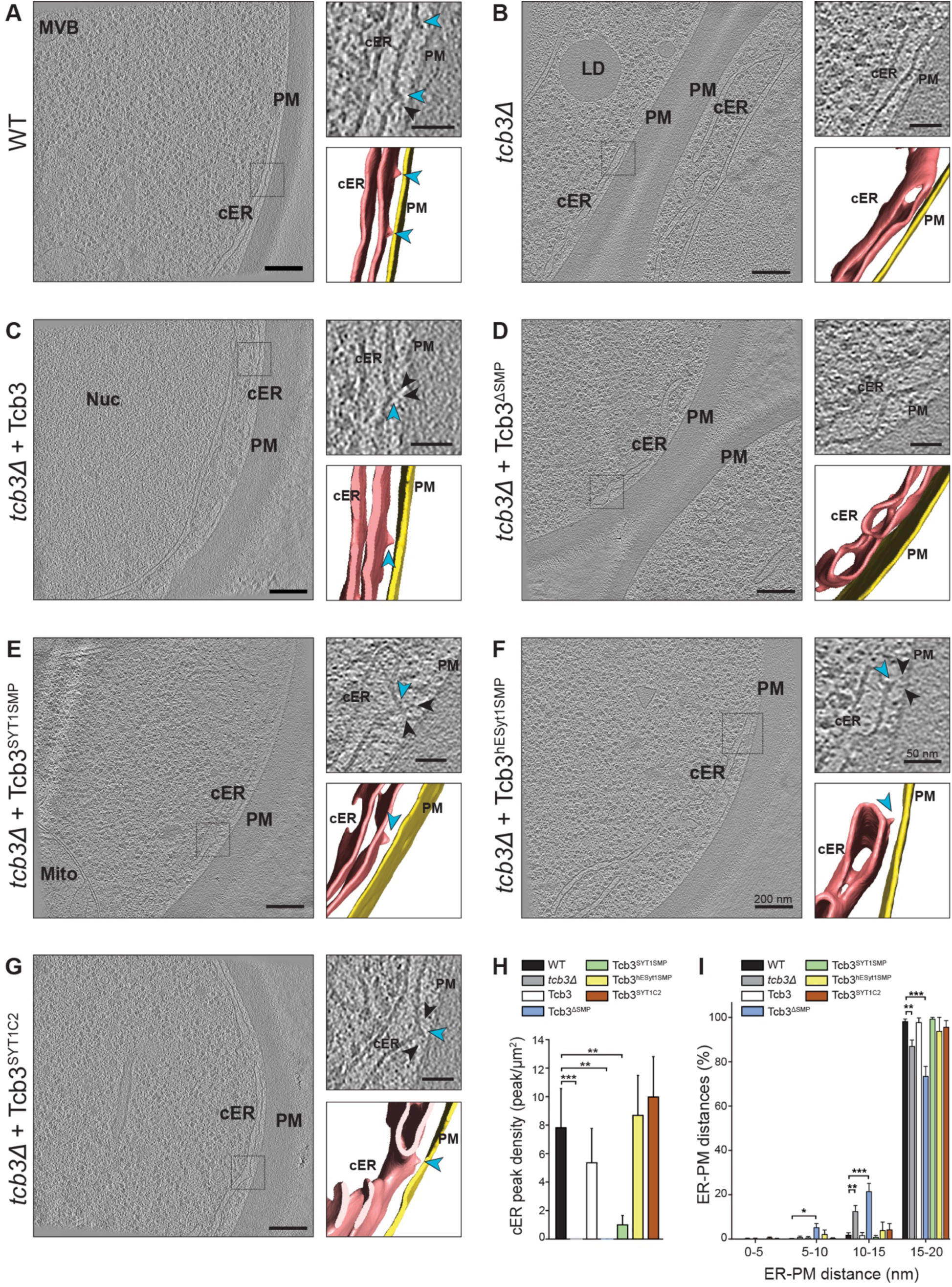
Cryo-ET of Tcb3-mediated ER-PM MCS in yeast. **A-G:** Cryo-ET of yeast cells of the indicated strains. Left: 1.2 nm-thick tomographic slice of yeast cells showing the cortical ER (cER), the plasma membrane (PM) and other cellular elements as indicated. Nuc: nucleus, Mito: mitochondrion, LD: lipid droplet. Right: enlargement of the boxed region (top) and corresponding 3D membrane segmentation (bottom). PM and cER are depicted in yellow and pink, respectively. Blue arrowheads point to cER peaks, black arrowheads point to densities connecting the cER peaks and the PM. **H:** cER peak density per µm^2^ of cER membrane area showing average values (gray bars) + SE (error bars). ** and *** indicate p-value < 0.01 and p-value <0.001, respectively using one-way Anova and Tuckey multiple comparison test. **I:** histogram distribution of ER-PM distances in indicated strains. *, ** and *** indicate p-value < 0.05, p-value < 0.01 and p-value < 0.001, respectively using two-way Anova and Tuckey multiple comparison test. For H and I: N = 16 (WT), 22 (*tcb3*Δ), 17 (*tcb3*Δ + Tcb3-GFP), 7 (*tcb3*Δ + Tcb3^ΔSMP^), 13 (*tcb3*Δ + Tcb3^SYT1SMP^), 10 (*tcb3*Δ + Tcb3^hE-Syt1SMP^) and 12 (*tcb3*Δ + Tcb3^SYT1C2^) ER-PM MCS (see Table S6). In H and I, significance is asserted in comparison to WT. Significance is only displayed for selected comparisons. P-values for all comparisons can be found in Tables S4 and S5.

To further characterize the morphology of the ER-PM MCS formed in the presence of these constructs, we systematically measured the distance between the ER and the PM for all conditions (Figure 5I). We found that the mean ER-PM distance was not significantly different from the WT for all strains analyzed (Table S6), suggesting that the Tcb3 mutants studied did not cause major reorganizations of the cER. However, in cells lacking either Tcb3 (*tcb3*Δ) or the Tcb3 SMP domain (Tcb3^ΔSMP^), ER-PM distances under 15 nm were significantly enriched compared to cells expressing SMP-containing constructs (Figure 5I, Table S5), including Tcb3^SYT1SMP^, which did not show significant functional rescue. These observations suggest that the E-Syt SMP domain acts as a “molecular ruler”, maintaining a defined ER-PM distance. Maintenance of ER-PM distance correlates with Tcb3 cER localization but not with stress tolerance, which is linked to cER peak formation.

## Discussion

In this study, we dissect the roles of the conserved E-Syt modules (N-terminal ER anchor, SMP and C2 domains) in driving E-Syt ER-PM MCS localization and stress tolerance. We demonstrate that targeting of E-Syts to the cER requires all three modules, and that exclusive cER localization is necessary but not sufficient for E-Syt function in stress-resistance. Additionally, our cryo-ET analyses of ER-PM MCS show that E-Syts act as rulers to maintain a defined ER-PM distance, and reinforce the direct relationship between E-Syt function in PM integrity maintenance and the formation of high-curvature membrane peaks at the cER^10^.

The Tcb3 unstructured N-terminal tail regulates Tcb3 mobility^49^, but appears dispensable for both the cER localization (Figure 1C) and the stress tolerance role of Tcb3 (Figure 1F). Also, the transmembrane domain is necessary but not sufficient to drive cER localization. Tcb3 and SYT1 both localize to tubular cER regions in yeast and plants, respectively^9–12^, though the Tcb3 transmembrane domain likely adopts a hairpin fold^9,50^, while SYT1 may use a single pass transmembrane domain ^29^ (Figure S1A). How both structural motifs bind tubular cER remains unclear, but our results show that they are not interchangeable (Figure 1C, E), in agreement with a previous report that substitution of the Tcb3 ER membrane anchor with the single transmembrane motif of Erg11 disrupted Tcb3 cER localization^12^. Thus, the requirements for cER localization appear different in yeast and plants. One possibility is that the membrane anchors recognize cER microdomains with different morphology and/or lipids, as yeast, plant and mammalian ER membranes have different lipid composition^51–53^. Another hypothesis is that the E-Syt N-terminal module engages in protein-protein interactions at ER-PM MCS. As SYT1 interacts with multiple reticulons at ER-PM MCS^54^, it is possible that in absence of plant reticulons, SYT1 N-terminus cannot localize at the yeast cER. In contrast, while direct binding has not been investigated, Tcb3 can localize to the cER in absence of reticulons^7,10^. However, the hairpin domain of Tcb3 could be involved in interactions with other proteins, which may not be possible for the single transmembrane domain of plant SYT1. Indeed, E-Syts can form homo- and heterodimers^9,19,22,27,28,50^ and the minimal domain required for dimerization of hE-Syt2 has been mapped to a segment encompassing the transmembrane domain^50^. Tcb3 requires either Tcb1 or Tcb2 for cER localization ^49,55^, and structural predictions suggest that their transmembrane hairpins are involved in Tcb2-Tcb3 heterodimerization^49^. Altogether, the E-Syt transmembrane segment may contribute to cER localization via interactions with both lipid and protein partners.

Our functional and cryo-ET analyses suggest that the mechanisms underlying PM targeting by C2 domains are conserved among E-Syts. We show that, in yeast, the last three C2 domains are sufficient to ensure Tcb3 function. Thus, while Tcb3 has two predicted Ca^2+^-binding C2 domains (C2C and C2D, Figure S1D), our results suggest that, similarly to other E-Syts, a single Ca^2+^-binding C2 domain might be sufficient. Furthermore, we demonstrate that the six C2 domains of Tcb3 can be functionally and structurally replaced by the two C2 domains of SYT1. Thus, localization (Figure 2B, D), PM integrity (Figure 2F) and cER peak formation (Figure 5G, H) can be performed by E-Syts containing a minimal set of C2s consisting of one Ca^2+^ binding domain (e.g. SYT1 C2A^30^) and one PM binding domain (e.g. SYT1 C2B^30^), which appear interchangeable among species. However, the additional C2 domains in yeast and metazoans might be required for fine regulation of E-Syt function and/or for its appropriate response to other stimuli^13,56^.

Our data aligns with previous studies showing that the E-Syt SMP domain is required for enrichment at the cER, possibly via E-Syt homo- or heterodimerization^9,10,19–22,28,49,55^. In this scenario, our findings that interspecies SMP domains enable ER-PM MCS localization suggest that their cross-dimerization is possible, consistent the predicted overall structural similarity between the SMP domain of Tcb3, hE-Syt1, SYT1 and dmE-Syt (Figure S1F), and hE-Syt2 (Figure S1E). Interestingly, we (Figure 5) and others^49^ find that E-Syt cER localization and correct ER-PM distance are perfectly correlated, suggesting that these two phenomena are mechanistically linked. SMP dimers may act as spacers, defining a minimum ER-PM distance of ∼15 nm. However, as SMP dimers form ∼9 nm-long rods^20^ (Figure S1E), additional elements may contribute to bridging the ER-PM gap.

Importantly, only some interspecies dimers may be functional stress responders. While predictions of SMP domains for Tcb3, hE-Syt1 and dmE-Syt show lipid-binding channels of similar width, SYT1 likely has a wider, uncapped channel due to the absence of helix α2 (Figure S1F), possibly altering lipid binding properties^13,20,21,57,58^. This divergence may impair lipid transport by yeast/metazoan heterodimers with plant SMPs, explaining why SYT1 chimeras carrying the Tcb3 or hE-Syt1 SMP domain can target to ER-PM MCS in plants (Figure 4C, Figure S3A), but fail to rescue the *syt1* defect in cold stress (Figure 4D, E). Equivalently, replacing the Tcb3 SMP with that SYT1 and hE-Syt1 yields WT-like cortical localization in yeast (Figure 3B, C), but only the human SMP can confer stress resistance (Figure 3D). Furthermore, only SMP dimers providing stress resistance give rise to cER peaks in yeast (Figure 5), supporting the hypothesis that E-Syt-mediated cER peaks preserve PM integrity under stress by facilitating ER-to-PM lipid transfer^10^. Other curvature-sensitive E-Syt modules, such as the hairpin ER anchor^59–61^ and the C2 domains^62^, may also contribute to cER peak formation and/or E-Syt localization at cER peaks.

Altogether, our findings indicate that E-Syts localize to ER-PM MCS through the concerted action of conserved modules (Figure 6A), which simultaneously 1) sense the membrane curvature of cER tubules, possibly via lipid or protein interactions (transmembrane segment), 2) form homo/heterodimers (SMP domain), and 3) mediate PM binding (at least one C2 domain). Thus, E-Syts act as coincidence detectors for ER-PM MCS targeting. Once localized to ER-PM MCS, E-Syts fulfill a structural role by imposing a defined ER-PM distance. However, ER-PM MCS localization is not sufficient for E-Syt activity during stress, which additionally requires (Figure 6B): 1) SMP (hetero)dimers that can promote cER peak formation in yeast, and possibly also in metazoans, 2) SMP (hetero)dimers with species-specific lipid transport capacity, and 3) C2 domains that may sense stress-induced increases in cytosolic Ca^2+^.

**Figure 6:**
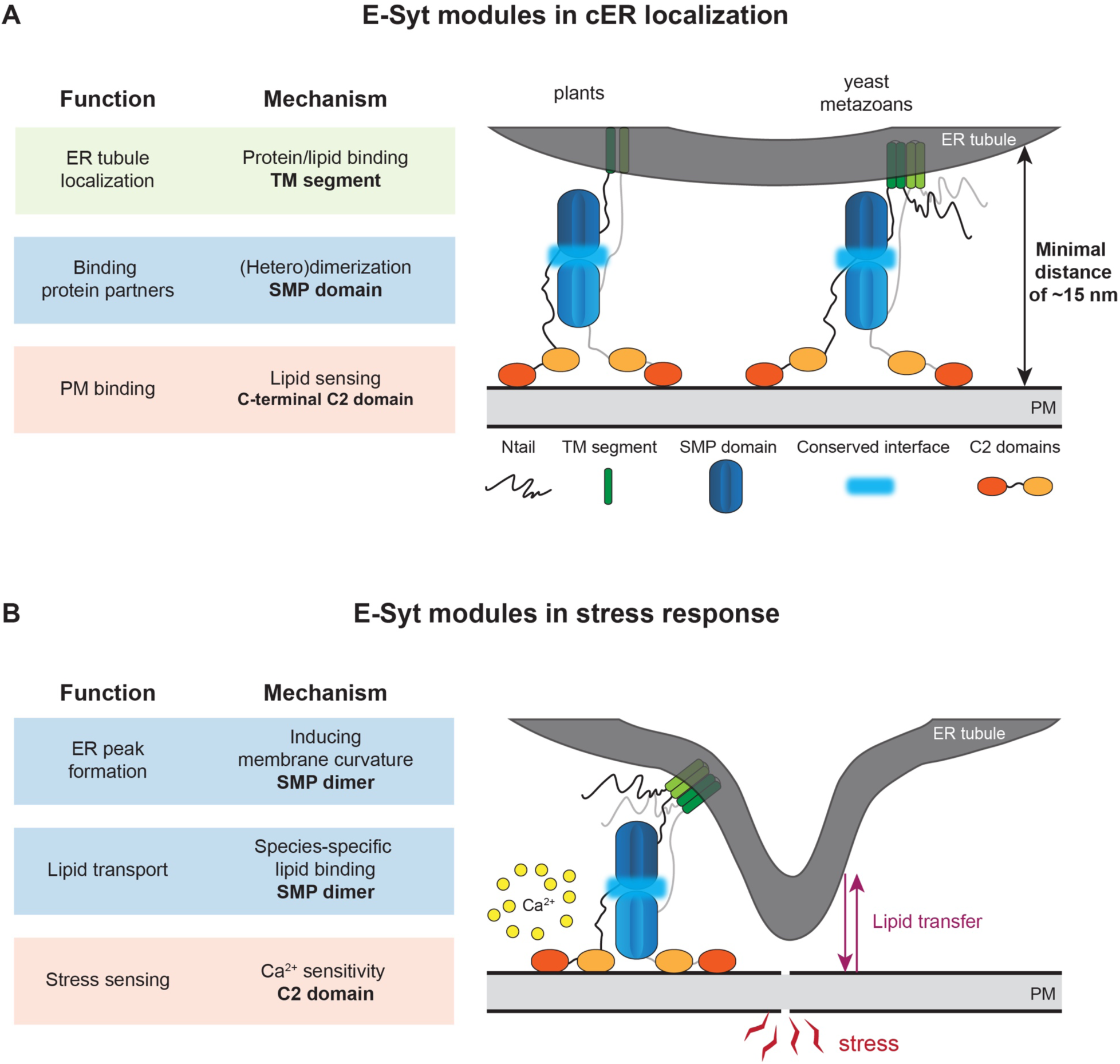
Model for the function of E-Syt domains at ER-PM MCS. This model summarizes the Discussion section, integrating findings of this study and the literature (see Discussion for references). **A**: E-Syts need the synergistic action of their three conserved modules for cER localization. The N terminal transmembrane segment (TM) targets E-Syts to tubular ER, possibly via lipid or protein interactions. The SMP forms E-Syt homo- and heterodimers through an interface that is sufficiently conserved to allow interspecies dimerization. The C2 domains are necessary to target E-Syts to the PM. Once properly localized to ER-PM MCS, E-Syts maintain a minimal ER-PM distance of ∼15 nm. **B**: cER localization of E-Syts is necessary but not sufficient for stress-response. This requires C2 domains for intracellular Ca^2+^ sensing and an SMP domain capable of functional dimerization for lipid transfer between the ER and the PM (not all heterodimers are functional). Functional SMP dimers also promote the formation of cER peaks, necessary for an efficient stress response possibly by facilitating intermembrane lipid exchange.

### Limitation of the study

This study provides a basis for understanding the molecular mechanisms governing E-Syt function under stress. Our work suggests that interspecies SMP dimerization is possible, but only some heterodimers can provide stress resistance. Future experiments should address the structural bases of SMP heterodimerization and lipid binding specificity, as well as the link between both phenomena. Further work is also needed to understand the precise mechanisms of stress sensing by E-Syts, including the roles of Ca^2+^. Lastly, deciphering the interplay between E-Syts and other parallel pathways (e.g. protein kinase C^63^) is essential to build an integrative picture of the cellular stress response.

## Resource availability

### Lead contact

Requests for further information and resources should be directed to and will be fulfilled by the lead contact, Rubén Fernández-Busnadiego.

## Materials availability

All unique reagents generated in this study are available from the lead contact with a completed materials transfer agreement.

## Data and code availability

The tomograms shown in Figure 5 are deposited in the Electron Microscopy Data bank (EMDB) under the following accession codes: EMD-57987, EMD-57988, EMD-57990, EMD-57991, EMD-57993, EMD-57994 and EMD-57995.

This paper does not report original code.

## Acknowledgements

We thank Dirk Schwitters for technical assistance, Tat Cheng for support with cryo-ET imaging, and Tanvir Shaikh for support with data analysis. We also acknowledge Eri Sakata and Alexander Neumann for their valuable suggestions, and all members of the Fernández-Busnadiego laboratory for helpful discussions. We thank Manuela Vega Sanchez, from the Servicio Central de Informatica (SCI), for technical assistance with the quantification of yeast confocal imaging, and Vitor Amorim-Silva for assistance with the molecular cloning. This work was supported by the Deutsche Forschungsgemeinschaft (DFG, German Research Foundation) under Germany’s Excellence Strategy (EXC 2067/1–390729940 Multiscale Bioimaging) and SFB1190 (project 22) to R.F-B. This work was also supported by PID2023-147983OB-100 awarded to M.A.B from the Spanish Ministry of Science and Innovation. This work was also supported by Grant PID2024-159647NB-I00 funded by MICIU/AEI/ 10.13039/501100011033 and by “ERDF A way of making Europe”, and the “European Union (ERDF/EU) to N.R-L. F.B-F was awarded with a Formación del Profesorado Universitario (FPU) fellowship (FPU17/03377) and a Ayuda de Movilidad para Estancias Breves fellowship (EST21/00714), from the Spanish Ministry for Universities. J.M-L was financed by the Ministry for Science and Innovation (Researcher Training Fellowship PRE2021-097655). R.P-M was financed by the Junta de Andalucía (Researcher Training Fellowship PREDOC_01435).

## Author contributions

J.C., F.B-F and J.K. constructed plasmids. J.C. and F.B-F performed plasma membrane integrity assays. F.B-F, J.M-L and R.P-M carried out plant experiments and confocal microscopy of yeast and plants. F.B-F and J.M-L performed the quantification of the fluorescence data. J.C. and J.K performed cryo-ET experiments. J.K. performed tomogram segmentation, computational and statistical analysis of the cryo-ET data. J.C., F.B-F, J.M.L, J.K., M.A.B and R.F.-B. designed the research. M.A.B, F.B-F, J.M-L, N.R-L, R.P-M and R.F.-B. acquired the funding. J.K. wrote the manuscript with contributions from R.F-B, F.B-F, J.M-L, R.P-M and M.A.B.

## STAR★Methods

### Key resources table

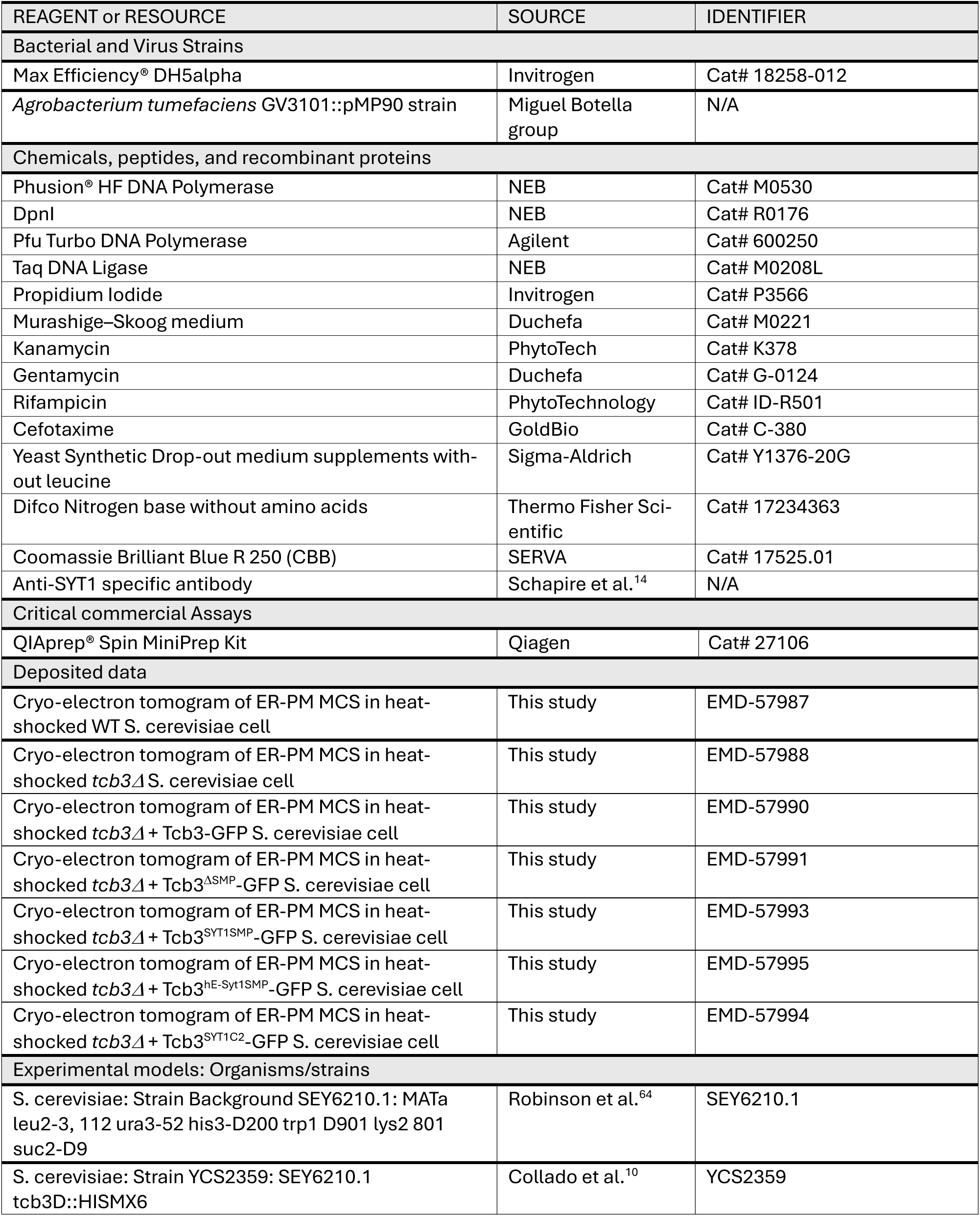
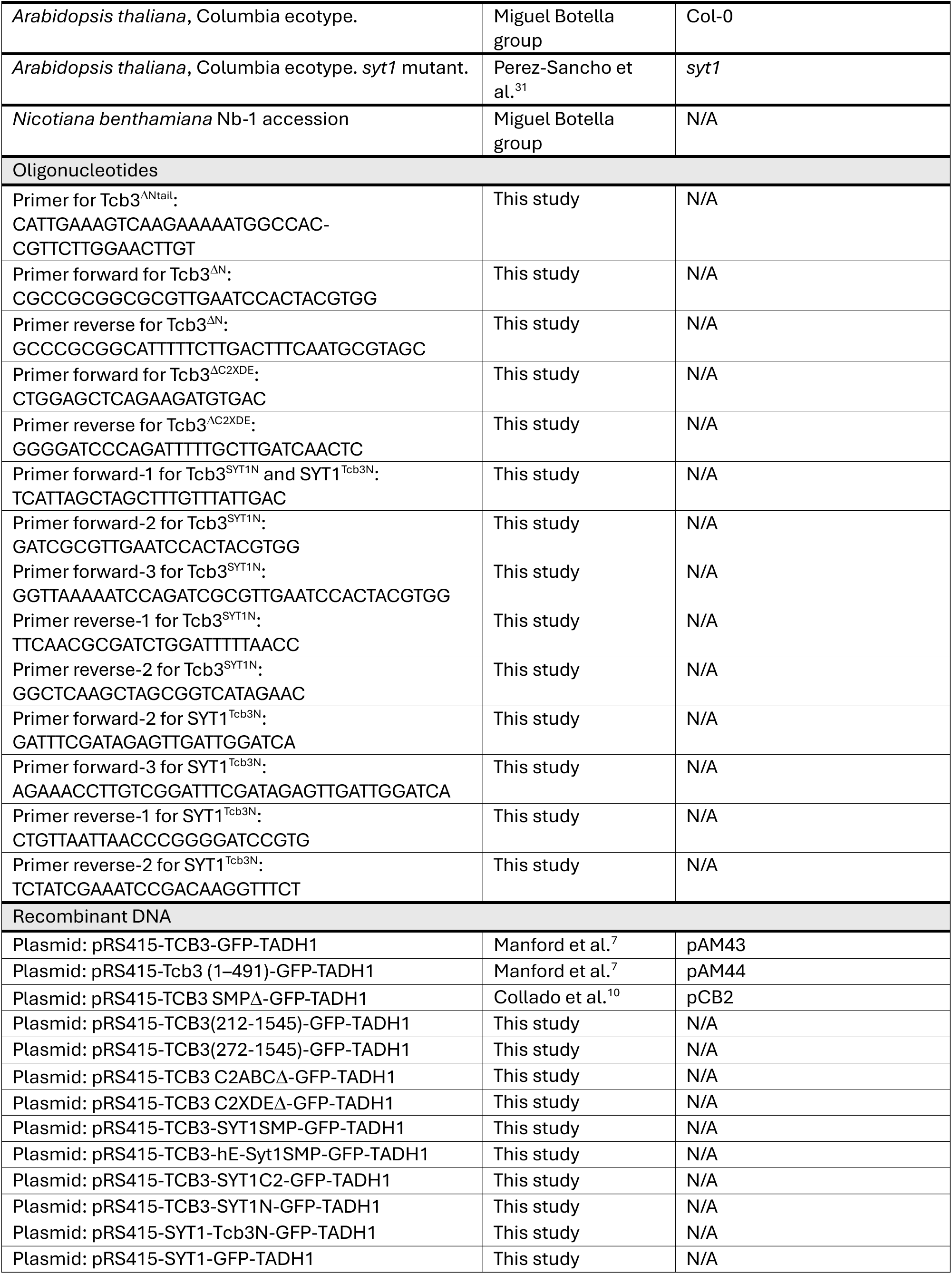
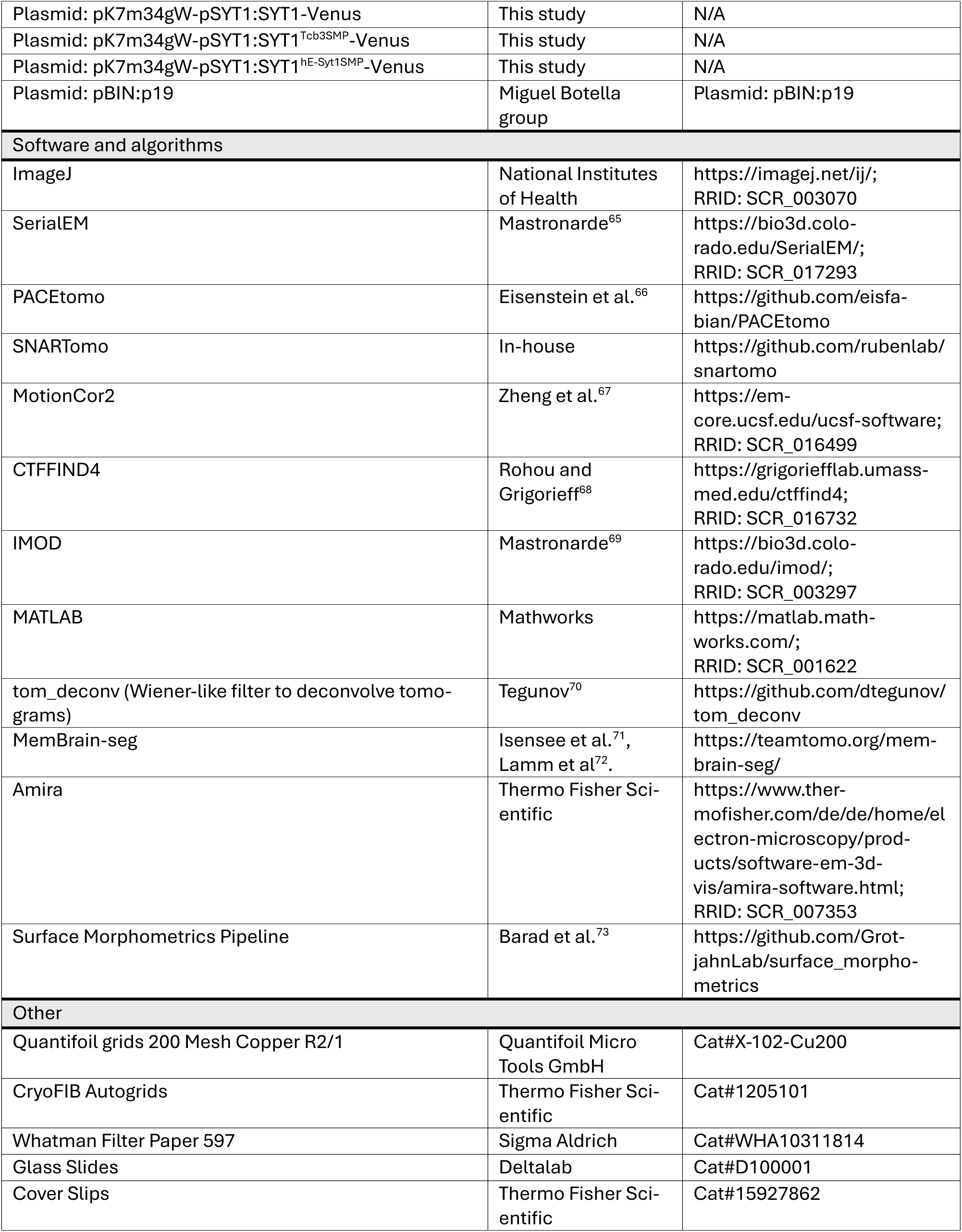

### Method details Plasmid construction

All Tcb3 deletions strains, except for Tcb3^ΔNtail^, Tcb3^ΔN^, Tcb3^ΔC2ABC^ and Tcb3^ΔC2XDE^ were previously reported^10^. Tcb3-SYT1and Tcb3-hE-Syt1 chimeras were created for this study. The Tcb3^ΔC2ABC^ mutant was generated from pAM43^7^ using restriction sites already present in the plasmid (SacII and NotI) and subsequent ligation using T4 ligase. The Tcb3^ΔN^ and Tcb3^ΔC2XDE^ mutants were generated by PCR from pAM43^7^ and subsequent ligation using T4 ligase. 1 µl of the ligation products were used to transform *E. coli* NEB 5-alpha strain. Tcb3^ΔNtail^ was constructed by deleting the first 211 amino acids of Tcb3 from pAM43^7^ using a single primer and one-step PCR. Briefly, 100 µM of the oligonucleotide was phosphorylated using 10U of T4 polynucleotide kinase (NEB) for 1 h at 37°C. The phosphorylated primer was then used for the PCR reaction, containing Pfu turbo polymerase (Agilent), Taq ligase (NEB) and pAM43 as a template. The PCR product was then digested with Dpn1 for 3 h at 37 °C. 5 µl of the digested PCR product was used to transform *E. coli* NEB 5-alpha strain. Tcb3^SYT1N^ and SYT1^Tcb3N^ were generated by PCR from pAM43^7^ and from the plasmid containing SYT1 full length (pRS415-SYT1-GFP-TADH1), followed by ligation using T4 ligase. 1 µl of the ligation products were used to transform *E. coli* NEB 5-alpha strain.

pSYT1:SYT1-Venus was generated using SYT1 CDS (AT2G20990)^22^ and the SYT1 promoter^43^ in triple Gateway ligations. The constructs Tcb3^SYT1SMP^, Tcb3^hE-Syt1SMP^ and Tcb3^SYT1C2^ for *S. cerevisiae* and SYT1^Tcb3SMP^ and SYT1^hE-Syt1SMP^ for *Arabidopsis* were generated by gene synthesis (GenScript) with codon optimization for expression in the corresponding organisms, adding restriction sites also present in pAM43^7^ for yeast constructs, or in pSYT1:SYT1-Venus, for *Arabidopsis* constructs. The chimeric constructs were then inserted into their respective destination plasmids using restriction enzymes, namely NheI and BamHI for yeast, and BamHI and XbaI for *Arabidopsis*.

### Yeast cell culture

WT cells (strain SEY6210.1^64^) were grown on plates containing synthetic complete medium, *tcb3*Δ yeast cells^10^ were grown on selection plates containing synthetic medium lacking histidine and *tcb3*Δ yeast carrying plasmids were first grown on selection plates containing synthetic media lacking leucine to maintain the plasmid selection for two days. All yeast strains were plated on their respective medium supplemented with 2% glucose. Yeast colonies were then inoculated in liquid selection media and incubated at 30 °C until reaching 0.8 OD_600_.

### Plant material

We used *Nicotiana benthamiana* Nb-1 accession (tobacco) for transient expression. For stable expression and cold-stress assays, we used *Arabidopsis thaliana*, ecotype Col-0, as wild-type, as well as the already characterized mutant line *syt1* (SAIL_775_A08)^14,31^.

### Generation of *Arabidopsis* stable cell lines

Constructs used in *Arabidopsis* were transformed into the *syt1* mutant line by floral dipping^74^. Once harvested, *Arabidopsis* seeds were sterilized by the chlorine gas method^75^ (100 ml bleach + 3 ml 37% HCl) and plated onto Murashige–Skoog (MS) media containing 50 μg/mL kanamycin and 300 μg/mL cefotaxime. Resistant first generation of transgenic (T1) plants were transferred to soil and grown to senescence. Collected seeds from the second generation of transgenic (T2) plants were grown on half strength MS (MS ½) plates without antibiotic for 7 days and used for construct expression screening by western blot, using an α-SYT1 antibody targeting SYT1 C2 domains^14^. Coomassie Brilliant Blue staining served as loading control. Parallelly, T2 seeds were also plated onto MS ½ media containing 50 μg/mL kanamycin for studying the transgene insertion number by the ratio of resistant:sensitive seedlings. Transgenic lines with 3:1 resistant:sensitive ratio, implying the presence of a single transgene insertion, were selected and used to obtain homozygous lines. Single insertion lines with high expression of the transgene were selected by western blot (Figure S3B) and used for confocal microscopy and cold-stress assays.

### Arabidopsis manipulation and growth conditions

*Arabidopsis* seeds were sterilized by chlorine gas. Seeds were plated on MS ½ with 1.5% (w/v) sucrose and 0.8% (w/v) agar. Plated seeds were vernalized for three days at 4 °C in darkness, then grew vertically on long-day photoperiod (16 h light/8 h dark; 120 µmol photons/m^2^·s, 22 °C).

### Transient expression in *Nicotiana benthamiana*

SYT1-GFP, MAPPER, SYT1^Tcb3SMP^ and SYT1^E-SYT1SMP^ were co-transformed with a plasmid carrying the viral p19 gene into *Agrobacterium tumefaciens* (GV3101::pMP90) by electroporation. The *A. tumefaciens* strains were grown in liquid LB media containing 50 μg/mL kanamycin, 25 μg/mL gentamycin and 50 μg/mL rifampicin overnight. Cultures were centrifuged (3000 g, 15 min), resuspended in infiltration solution (10 mM MES pH 5.6, 10 mM MgCl2, 1 mM acetosyringone) and incubated 2 h in the dark. The third and fourth leaves from the apex of 3-week-old plants, were infiltrated from the abaxial side with a needleless syringe. Plants were grown for two more days prior to confocal microscopy.

### Cold treatment for cell viability in *Arabidopsis thaliana*

The experiments were performed as previously described^19^. Briefly, 6-day-old *Arabidopsis* seedlings were put into ice-cold (4 °C) MS 1/10 liquid media for 30 min, then transferred to MS 1/10 room temperature MS liquid media with 10 μg/mL fluorescein diacetate (FDA, Sigma Aldrich) for 5 min. The seedlings were washed at room temperature with MS 1/10 liquid media by gentle agitation and immediately used for imaging.

### Confocal microscopy imaging

Yeast cells were grown on solid minimum medium (0.17% (w/v) yeast nitrogen base, 0.5% (w/v) ammonium sulphate, 2% (w/v) glucose, pH 6.5, and 2% (w/v) agar for solid medium) supplemented with amino acids, at 30 °C for two days. Colonies were grown overnight in minimum liquid media supplemented with amino acids, and 10 µL were transferred to a new liquid media tube in the morning. After 5 h, cells in exponential phase were directly visualized on glass slides with the confocal microscope.

For tobacco, leaf disks were cut from the plant leaves two days post infiltration, and immediately before visualization. For *Arabidopsis*, 3-days-old cotyledons were used for confocal analysis.

Confocal images for subcellular localization studies in yeast and plants were taken using a LSM880 confocal microscope (Zeiss), equipped with a 458-nm argon laser for GFP and Venus excitation and a GaAsP (*Arabidopsis*) or a PMT (tobacco and yeast) detector. A Plan-Apochromat 63x/1.40 Oil DIC M27 objective was used, with a 2x (*Arabidopsis* and tobacco) or

3x (yeast) digital zoom. Selected regions were cropped for the figures. Confocal images of *Arabidopsis* seedlings stained with FDA in cold stress experiments were taken using a Stellaris 8 confocal microscope with a HC PL APO CS2 20x/0.75 IMM objective (Leica), a WLL 491nm laser, a PMT detector and a x0.75 digital zoom. All image processing was done using FIJI^76^.

### PM integrity assays

Yeast cells were grown at 26 °C to mid-log phase in synthetic complete medium or synthetic selective medium as required. For heat shock, cells were shifted to 42 °C for 10 min, resuspended in PBS, and incubated with 25 µg/ml propidium iodide (Invitrogen) for 10 min. Cells were then washed twice with ddH2O and analyzed by flow cytometry (CytoFLEX V2-B2-R0 Flow Cytometer). For each condition, 10,000 cells were measured and the background was determined by analyzing the cells prior to staining with propidium iodide.

### Yeast cell vitrification

Cryo-EM grids (R2/1, Cu 200 mesh grid, Quantifoil) were glow discharged using a plasma cleaner (PDC-3XG, Harrick) for 30 s and mounted on a Vitrobot Mark IV (Thermo Fischer Scientific). Yeast cells were incubated at 42°C for 10 min and a 3.5 µl drop of the yeast culture was deposited on the carbon side of the EM grid and blotted from the back to remove excess liquid using filter paper (Whatman 597). Grids were then immediately plunged frozen into a liquid ethane/propane mixture at liquid nitrogen temperature and stored in grid boxes immersed in liquid nitrogen until further need.

### Cryo-focused ion beam milling

Vitrified EM grids were mounted into Autogrid carriers (Thermo Fischer Scientific), held in place by a copper ring. An Aquilos 2 cryo-focused ion beam (FIB)/scanning electron microscope (SEM) (Thermo Fisher Scientific) was used to prepare lamellae. A protective layer of organometallic platinum was deposited on the grid with the gas injection system for 40 seconds. The sample was tilted to an angle of 20° and < 200 nm thick lamellae were prepared in two steps: first, using the Ga^2+^ ion beam at 30 kV and 300 pA beam current for rough milling (until ∼600 nm thick lamella) and then by fine milling at 30 kV and 50 pA. The milling process was monitored using SEM imaging at 3 kV and 13 pA.

### Cryo-electron tomography

Data acquisition: A Krios G4 Cryo-transmission electron microscope (Thermo Fisher Scientific) equipped with a 300 kV field emission gun, Selectris energy filter, and a Falcon 4i direct electron detector camera (Thermo Fisher Scientific), was used for data collection. Tilt series of the lamellae were acquired using PACE-tomo ^66^ implemented within SerialEM 4.0^65^ at a magnification of 33000x (3.653 Å/pixel) and a defocus of -5 to -7 µm, using a dose-symmetric acquisition scheme^77^ from -45° to +63° at increments of 3°, and a target total dose per tomogram of around 120 e-/A^2^.

Tomogram reconstruction: Tilt series preprocessing was automated using an inhouse script (https://github.com/rubenlab/snartomo) performing frame alignment using Motioncor2^67^, CTF estimation using CTFFIND 4.1.14^68^, tilts series alignment using patch tracking and weighted back-projection for tomogram reconstruction using IMOD 4.11^69^. Tomograms were binned twice to 14.61 Å/pixel and filtered using the Wiener-like filter tom-deconv^70^.

Tomogram segmentation: automatic membrane segmentation was performed using Mem-Brain-Seg^71,72^. The segmentation was manually corrected and colored using Amira (Thermo Fisher Scientific).

### Quantification and statistical analysis

PM integrity assays: Average values from at least four independent biological repeats were analyzed for all conditions. Bar plots (Figure 1-3) show mean values for each condition + SEM. Statistical significance was assessed using two-way Anova and Tukey’s multiple comparison test. See also Table S1.

Subcellular localization: cER/non-cER ratios: Confocal images of the equatorial plane of yeast cells in exponential phase were processed in FIJI^76^ and the fluorescent signal was quantified using a macro as follows: fluorescence images (Figure S2A) were segmented using the Threshold module, creating a mask of the cells. Touching cells were split using Watershed segmentation and contours of the cells were detected using the Particle Analyzer module (Figure S2B). Contours were manually examined to remove artifacts (Figure S2B). Curated contours defined the cell outlines (Figure S2C). Cell outlines were then shrunk by 10 pixels (Figure S2C). The region between the two outlines defined the cER region (Figure S2D) while the region within the second outline defined the non-cER region (Figure S2D). Measurements of the fluorescence intensity (mean pixel intensity) were normalized by area, and a ratio of cER/non-cER signal was obtained per cell. Bar plots (Figure 1-3) show the mean ratios of cER/non-cER + SE for each condition. Statistical significance was assessed using one-way ANOVA and Tukey’s multiple comparison test. See also Table S2.

Cold stress experiments: processing of confocal images of *Arabidopsis* seedlings stained with FDA was performed using FIJI^76^ as follows: Z-projections from seedling roots were converted into binary images with the Auto Threshold module using the Moments algorithm ^78^. As FDA is only fluorescent in living cells, binarized images show living cells in white and dead cells in black. Five different regions of interest (ROIs, size 73.48 x 73.48 μm) were chosen for each root, and the area of the white signal was measured. The ROI corresponding to the root apical meristem was not analyzed but used as reference. In Figure 4E, the data was expressed as percentage of total area by dividing the white signal by the total ROI area. Statistical significance was assessed using one-way Anova followed by Tukey’s multiple comparison test. See also Table S3.

cER peak density: This represents the number of ER peaks per cER surface. cER surface was calculated as the sum of the area of all cER triangles divided by two, to only take in account the ER surface facing the PM. The bar plot in Figure 5H shows the mean value + SE of peak density per condition. Statistical significance was estimated using one-way Anova followed by Tukey’s multiple comparison test. See also Table S4.

ER-PM distance measurements: The distance between ER and PM was calculated using the surface morphometrics pipeline^73^. Briefly, voxel-based segmentations were converted to surface meshes using the screened Poisson algorithm^79^, which were then simplified into triangle graphs. The nearest distance to the closest triangle in the PM was measured for each triangle in the ER membrane using a k-dimensional tree. As triangles belonging to the same surface are not independent measurements, a generalized linear mixed model was fitted to the data for quantification. Distance distributions were represented as a histogram (Figure 5I) and statistical significance was assessed by two-way Anova and Tukey’s multiple comparison test. See also Tables S5 and S6.

## Supplemental information

Figures S1-S3, Tables S1-S6.

**Figure S1:**
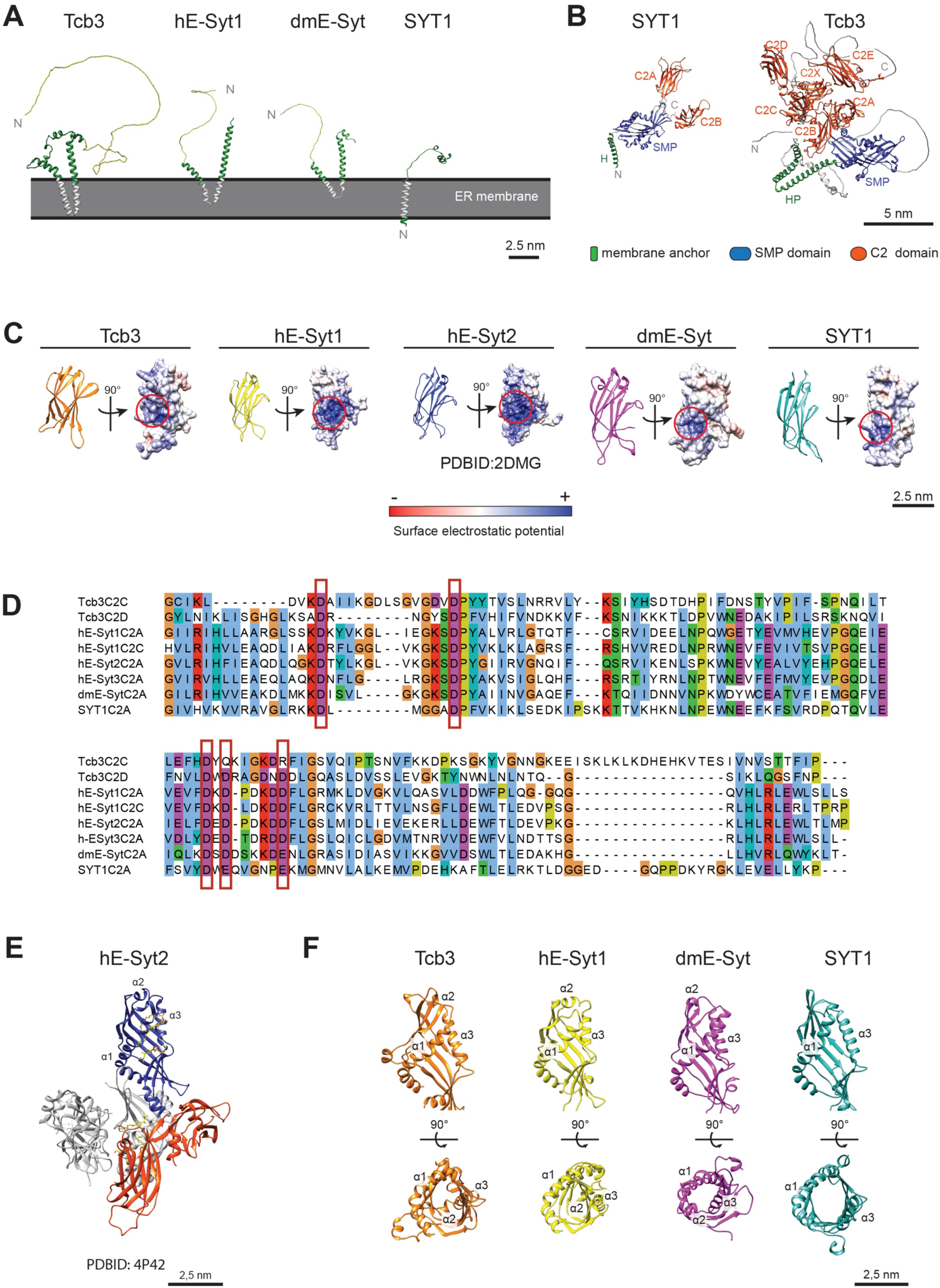
Structural features of E-Syts. **A:** Mapping of hydrophobic residues (light grey) on AlphaFold2 structural predictions of the N-terminal transmembrane segment (green) of *Saccharomyces cerevisiae* Tcb3, *Homo sapiens* hE-Syt1, *Drosophila melanogaster* dmE-Syt and *Arabidopsis thaliana* SYT1 suggests how E-Syts could be anchored to the ER membrane (grey). The N-terminal unstructured tail of *Saccharomyces cerevisiae* Tcb3, *Homo sapiens* hE-Syt1, *Drosophila melanogaster* dmE-Syt is depicted in light green. **B:** AlphaFold2 structural predictions of full-length *Arabidopsis thaliana* SYT1 and *Saccharomyces cerevisiae* Tcb3. N, N-terminus, C, C-terminus, HP, hairpin domain, H, helix. **C:** Electrostatic surface analysis of AlphaFold2 structural predictions of the C-terminal C2 domain of Tcb3, hE-Syt1, dmE-Syt and SYT1 and the experimental structure of hE-Syt2 C-terminal C2 domain (PDBID: 2DMG). Red circles indicate positively charged residues at equivalent positions. **D**: Sequence alignment of predicted Ca^2+^-binding C2 domains of Tcb3, h-E-Syt1-3, dmE-Syt and SYT1. Red rectangles indicate the position of the canonical Ca^2+^ residues of C2 domains. **E:** Crystal structure of the SMP domain, C2A and C2B domains of ESyt2 (PDBID: 4P42). The protein construct dimerizes through the SMP domain. One monomer is colored in grey while the other is colored as in A. **F:** AlphaFold2 structural prediction of E-Syts SMP domains. α1, α2, α3 label the α-helices of the SMP domains.

**Figure S2:**
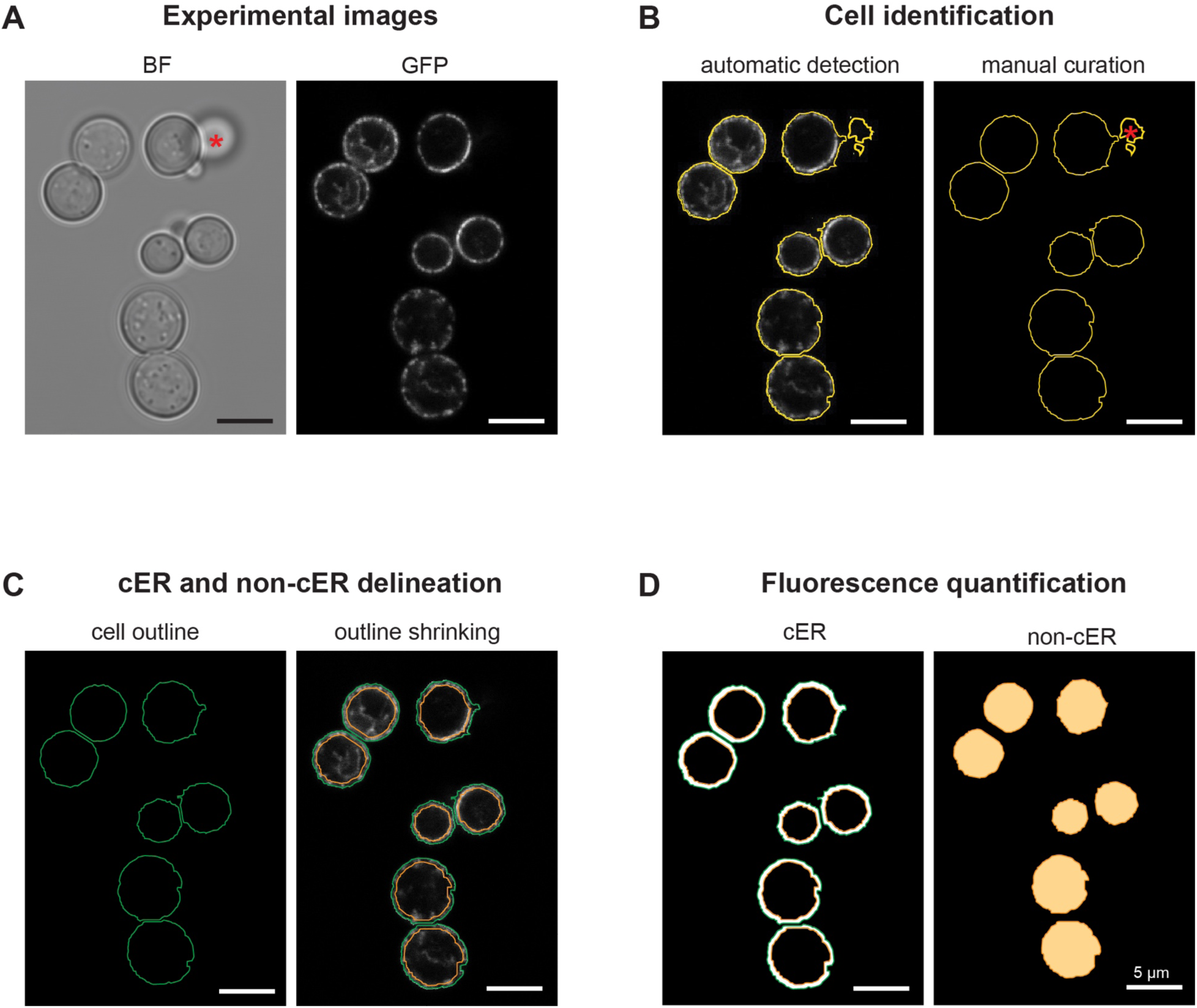
Quantification of subcellular localization of Tcb3-GFP constructs in yeast. The fluorescence was quantified using a FIJI^76^ macro implementing the following steps. **A**: confocal images of the equatorial plane of yeast cells in exponential phase were used as input. BF: bright field. **B**: Fluorescence images were segmented and cell outlines (yellow contours) were automatically detected using the Particle analyzer module, and manually examined to remove artifacts (red stars). **C**: Curated cell outlines (green contours) were subsequently shrunken to delineate the non-cER region of the cells (orange contours). **D**: The fluorescence signal (mean pixel intensity) quantified in the region between the outline of the cell (green contour) and the non-cER region (orange contour) was assigned to the cER (white surface), while the fluorescence signal within the orange contour (orange surface) was assigned to the non-cER region. The values were subsequently normalized by area.

**Figure S3:**
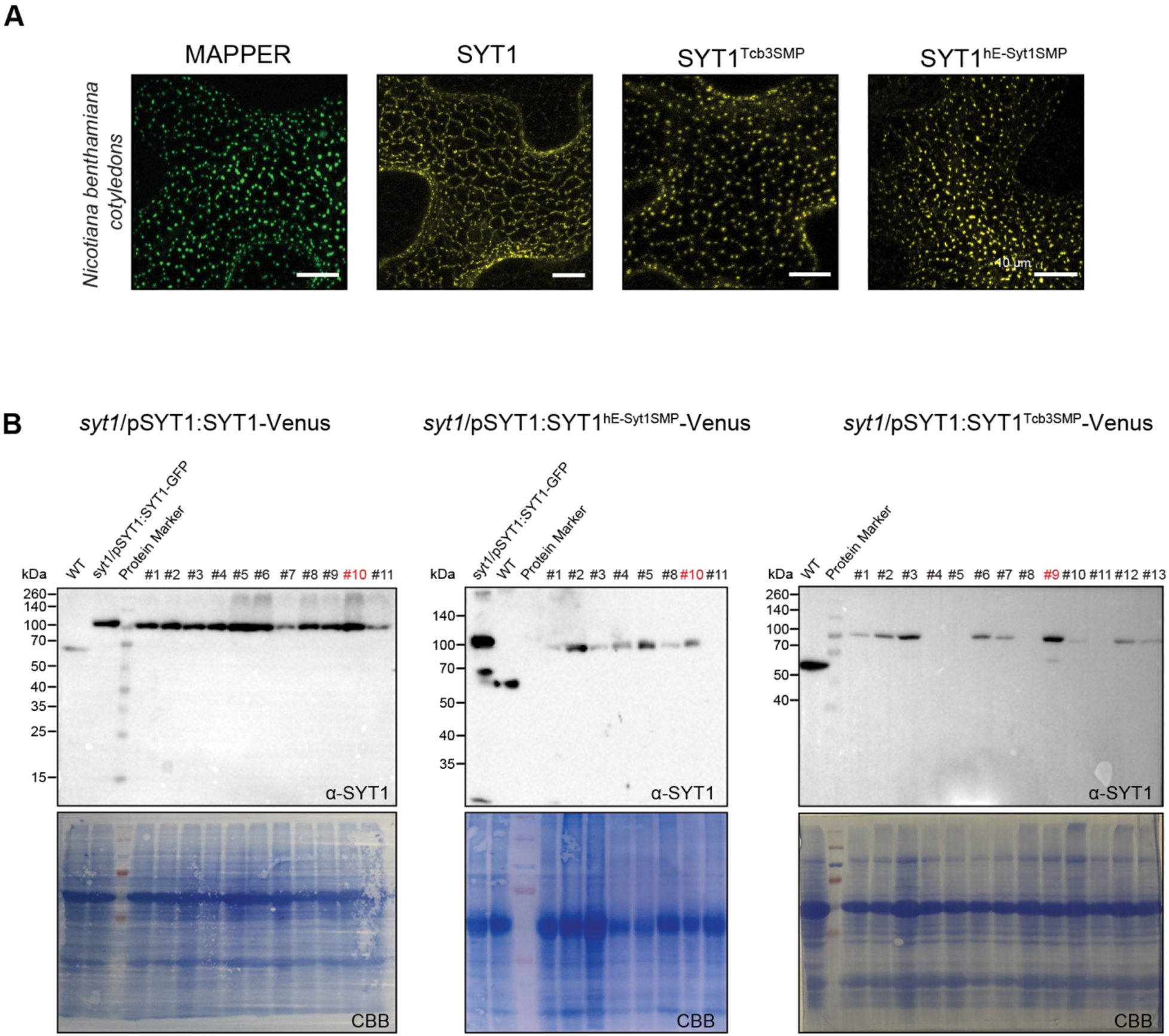
Transient expression and subcellular localization of the E-Syt SMP domain in *Nicotiana benthamiana* (tobacco) and line selection in *Arabidopsis*. **A:** Subcellular localization of SYT1, MAPPER and SMP constructs at the cortical region of epidermal cells of transiently transformed tobacco plants by confocal imaging. **B:** Western blots for the selection of T1 Arabidopsis stable lines. Upper panels, α-SYT1 antibody targeting SYT1 C2 domains. Lower panels, Coomassie Brilliant Blue staining (CBB) as loading control. Stable lines indicated in red were selected for further experiments. WT indicates endogenous SYT1. *syt1*/pSYT1:SYT1-GFP serves as a control to follow the expression of SYT1-Venus, as GFP and Venus have similar sizes.

**Table S1:** Statistics of PM integrity assays related to Figures 1-3. The p-values compare the percentage of cells with compromised PM integrity at 42°C using two-way Anova followed by Tukey’s multiple comparison tests.

| Figure 1 | Reference | Measurement | p-value |
| --- | --- | --- | --- |
|  | WT | *tcb3Δ* | < 0.0001 |
|  | WT | *tcb3Δ* + Tcb3 | > 0.9999 |
|  | WT | *tcb3Δ* + Tcb3<sup>ΔNtail</sup> | > 0.9999 |
|  | WT | *tcb3Δ* + Tcb3<sup>ΔN</sup> | < 0.0001 |
|  | WT | *tcb3Δ* + SYT1 | < 0.0001 |
|  | WT | *tcb3Δ* + Tcb3<sup>SYT1N</sup> | < 0.0001 |
|  | WT | *tcb3Δ* + SYT1<sup>Tcb3N</sup> | < 0.0001 |
|  | *tcb3Δ* | *tcb3Δ* + SYT1<sup>Tcb3N</sup> | 0.0250 |
|  | *tcb3Δ* + SYT1 | *tcb3Δ* + SYT1<sup>Tcb3N</sup> | 0.0078 |
| Figure 2 | WT | *tcb3Δ* | <0.001 |
|  | WT | *tcb3Δ* + Tcb3 | 0.9993 |
|  | WT | *tcb3Δ* + Tcb3<sup>ΔC2</sup> | <0.01 |
|  | WT | *tcb3Δ* + Tcb3<sup>ΔC2ABC</sup> | <0.001 |
|  | WT | *tcb3Δ* + Tcb3<sup>ΔC2XDE</sup> | <0.001 |
|  | WT | *tcb3Δ* + SYT1 | < 0.0001 |
|  | WT | *tcb3Δ* + Tcb3<sup>SYT1C2</sup> | > 0.9999 |
| Figure 3 | WT | *tcb3Δ* | < 0.0001 |
|  | WT | *tcb3Δ* + Tcb3 | > 0.9999 |
|  | WT | *tcb3Δ* + Tcb3<sup>ΔSMP</sup> | < 0.0001 |
|  | WT | *tcb3Δ* + SYT1 | < 0.0001 |
|  | WT | *tcb3Δ* + Tcb3<sup>SYT1SMP</sup> | <0.001 |
|  | WT | *tcb3Δ* + Tcb3<sup>hE-Syt1SMP</sup> | 0.9637 |
|  | *tcb3Δ* | *tcb3Δ* + Tcb3<sup>SYT1SMP</sup> | > 0.9999 |

**Table S2:** Statistics of cER localization related to Figures 1-3. The p-values compare the average ratio between the cortical and non-cortical fluorescence using one-way Anova followed by Tukey’s multiple comparison tests.

| Figure 1 | Reference | Measurement | p-value |
| --- | --- | --- | --- |
|  | *tcb3Δ* + Tcb3 | *tcb3Δ* + Tcb3<sup>ΔN</sup> | <0.001 |
|  | *tcb3Δ* + Tcb3 | *tcb3Δ* + Tcb3<sup>ΔNtail</sup> | 0.191 |
|  | *tcb3Δ* + Tcb3 | *tcb3Δ* + SYT1 | <0.001 |
|  | *tcb3Δ* + Tcb3 | *tcb3Δ* + Tcb3<sup>SYT1N</sup> | <0.001 |
|  | *tcb3Δ* + Tcb3 | *tcb3Δ* + SYT1<sup>Tcb3N</sup> | <0.001 |
| Figure 2 | *tcb3Δ* + Tcb3 | *tcb3Δ* + Tcb3<sup>ΔC2ABC</sup> | 0.603 |
|  | *tcb3Δ* + Tcb3 | *tcb3Δ* + Tcb3<sup>ΔC2XDE</sup> | 0.001 |
|  | *tcb3Δ* + Tcb3 | *tcb3Δ* + Tcb3<sup>SYT1C2</sup> | 0.274 |
| Figure 3 | *tcb3Δ* + Tcb3 | *tcb3Δ* + Tcb3<sup>ΔSMP</sup> | <0.001 |
|  | *tcb3Δ* + Tcb3 | *tcb3Δ* + Tcb3<sup>SYT1SMP</sup> | 0.941 |
|  | *tcb3Δ* + Tcb3 | *tcb3Δ* + Tcb3<sup>hE-Syt1SMP</sup> | 0.356 |

**Table S3:** Statistics of FDA staining related to Figure 4. The p-values compare the percentage of stained (alive) cells of different lines using one-way Anova followed by Tukey’s multiple comparison test.

| Reference | Measurement | p-value |
| --- | --- | --- |
| WT | *syt1* | <0.0001 |
| WT | *syt1* + SYT1 | > 0.9999 |
| WT | *syt1* + SYT1<sup>Tcb3SMP</sup> | <0.0001 |
| WT | *syt1* + SYT1<sup>hE-Syt1SMP</sup> | <0.0001 |

**Table S4:**
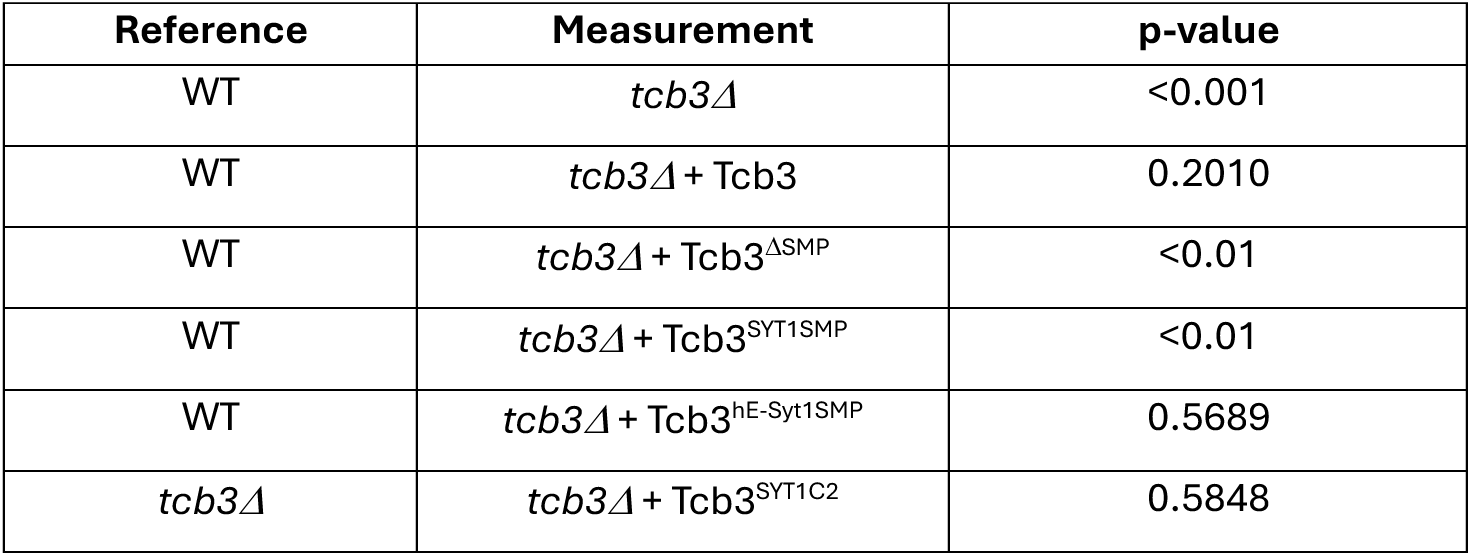
Statistics of cER peak density measurements related to Figure 5. The p-values compare the number of cER peaks per µm^2^ of ER using one-way Anova followed by Tukey’s multiple comparison test.

| Reference | Measurement | p-value |
| --- | --- | --- |
| WT | *tcb3Δ* | <0.001 |
| WT | *tcb3Δ* + Tcb3 | 0.2010 |
| WT | *tcb3Δ* + Tcb3<sup>ΔSMP</sup> | <0.01 |
| WT | *tcb3Δ* + Tcb3<sup>SYT1SMP</sup> | <0.01 |
| WT | *tcb3Δ* + Tcb3<sup>hE-Syt1SMP</sup> | 0.5689 |
| *tcb3Δ* | *tcb3Δ* + Tcb3<sup>SYT1C2</sup> | 0.5848 |

**Table S5:** Statistics of ER-PM distance measurements related to Figure 5. The p-values compare the mean ER-PM distance per bin using two-way Anova test and Tuckey multiple comparison test.

| Reference | Measurement | p-value<br>(0-5nm) | p-value<br>(5-10nm) | p-value<br>(10-15 nm) | p-value<br>(15-20 nm) |
| --- | --- | --- | --- | --- | --- |
| WT | *tcb3Δ* | NaN | 0.3430 | <0.01 | 0.0014 |
| WT | *tcb3Δ* + Tcb3 | 0.3370 | 0.4855 | 0.9288 | 0.8514 |
| WT | *tcb3Δ* + Tcb3<sup>ΔSMP</sup> | 0.1972 | 0.0307 | <0.001 | 0.0008 |
| WT | *tcb3Δ* + Tcb3<sup>SYT1SMP</sup> | NaN | 0.3332 | 0.504 | 0.4533 |
| WT | *tcb3Δ* + Tcb3<sup>hE-Syt1SMP</sup> | 0.4226 | 0.4515 | 0.6465 | 0.5578 |
| *tcb3Δ* | *tcb3Δ* + Tcb3<sup>SYT1C2</sup> | NaN | 0.6445 | 0.4435 | 0.4389 |

**Table S6:** Statistics of cryo-ET experiments related to Figure 5. N: number of experiments. “Total” indicates cumulated values for all tomograms analyzed in each condition. The total number of ER-PM distance measurements reflects the number of triangles from the ER membrane from which the distance to the closest triangle from the PM was measured. For ER-PM distance calculations, a generalized linear mixed model was fitted to the data to take in account that triangles belonging to the same membrane are not independent measurements. The total cER area analyzed is the sum of the area of all cER membrane triangles. The mean ER-PM distance calculation only took into account the distances measured within 35 nm. P-Values compare the mean ER-PM distance to WT using two-way Anova test and Tuckey multiple comparison test.

| Condition | N | Number<br>of ER-PM<br>MCS | Total number of<br>ER-PM distance<br>measurements | Total cER area<br>analyzed<br>($\mu\text{m}^2$) | Mean ER-PM<br>distance (nm) | p-value |
| --- | --- | --- | --- | --- | --- | --- |
| WT | 2 | 16 | 1081747 | 4.5 | 27.15 ± 0.24 | N/A |
| *tcb3Δ* | 2 | 22 | 937282 | 3.6 | 27.93 ± 0.20 | 0.10 |
| *tcb3Δ* + Tcb3 | 2 | 17 | 1117234 | 4.0 | 28.02 ± 0.44 | 0.60 |
| *tcb3Δ* + Tcb3<sup>ΔSMP</sup> | 2 | 7 | 1125233 | 3.8 | 26.80 ± 0.44 | 1 |
| *tcb3Δ* + Tcb3<sup>SYT1SMP</sup> | 2 | 13 | 1178365 | 4.9 | 28.45 ± 0.40 | 0.08 |
| *tcb3Δ* + Tcb3<sup>hE-Syt1SMP</sup> | 2 | 10 | 937731 | 3.1 | 28.08 ± 0.37 | 0.06 |
| *tcb3Δ* + Tcb3<sup>SYT1C2</sup> | 2 | 9 | 1187080 | 3.8 | 28.69 ± 0.60 | 0.24 |

